# Identification of a Neuroimmune Circuit that Regulates Allergic Inflammation in the Esophagus

**DOI:** 10.1101/2024.11.16.623883

**Authors:** Kendall Kellerman, Mia Natale, Eddie Gerstner, Yrina Rochman, Mark Rochman, Michael P. Jankowski, Marc E. Rothenberg

## Abstract

Eosinophilic esophagitis is a chronic food antigen-driven allergic inflammatory disease associated with symptoms involving the nervous system such as refractory pain. Yet, the role of the nervous system in disease pathogenesis has not received much attention. Herein, we demonstrate that allergen exposure evokes pain-like behavior in association with increased nociceptor signaling and transcriptional responses in dorsal root ganglia. NaV1.8+ sensory nerves were found traveling along the length of the esophagus, organized in distinct bundles adjacent to the basal epithelium, with beta III-tubulin+ sensory nerves distributed more distal to the lumen. Targeted deletion of *Il4ra* in NaV1.8+ neurons impeded allergen-induced increases in nerve innervation density. Furthermore, *Il4ra-/-^NaV1.8^ mice* had diminished allergen-induced allergic inflammation in the esophagus including eosinophilia and transcription of pro-inflammatory genes. Translational studies revealed extensive myelinated nerve innervation in the human esophagus, which was increased in patients with eosinophilic esophagitis. Taken together, these data indicate that allergic inflammation is associated with an increase in non-evoked pain, esophageal nerve density, altered sensitivity of sensory neurons, and transcriptional changes in dorsal root ganglia. These finding identify a type 2 neuroimmune circuit that involves the interplay of allergen-induced IL-4 receptor-dependent DRG responses that modify esophageal end-organ inflammatory responses.

## Introduction

Eosinophilic esophagitis (EoE) is a chronic allergic inflammatory disease often accompanied by refractory symptoms such as pain, suggesting the interplay of the neurological and immunological systems. While esophageal eosinophilia is a hallmark of the disease, there is often a dissociation between eosinophil levels and symptoms (1–3). For example, patients in histological remission (i.e. <15 eosinophils/hpf) often have persistent symptoms (2, 3). In fact, eosinophil depleting antibodies eliminate eosinophils but do not improve symptoms in eosinophilic gastrointestinal diseases (4, 5). Taken together, the mechanism of symptom development in EoE is likely more complex than eosinophilic inflammation alone. Indeed, EoE pathogenesis is mediated by the interplay of the innate and adaptive immune system, the latter driven by food antigen-induced type 2 helper T cells, and their associated type 2 cytokines IL-4, IL-5 and IL-13. Recent studies demonstrate that the function of afferent nerves extends beyond their role in sensing peripheral stimuli including nociceptive signals, as neuropeptides released from sensory neurons can activate local immune cells (6–9). Conversely, sensory neurons can directly respond to type 2 cytokines, including IL-4, to induce itch and other allergic symptoms (6, 9). Taken together, evidence is emerging that the peripheral nervous system may contribute to the pathophysiology of allergic diseases, such as EoE.

A preliminary description of the nervous system involved in afferent sensing in the esophagus has been reported (9, 10). Afferent innervation of the esophagus consists of sensory neurons from both the spinal and vagal nerves (11, 12). The majority of esophageal spinal afferents converge with somatic input in the spinal cord. The corresponding somatic receptive fields are then distributed over the chest and forearm area (11), likely explaining symptoms of referred chest pain that EoE patients experience (13). Additionally, sensory neurons in the vagal nerve exhibit expression of canonical nociceptive ion channels comparable to that observed in DRG-derived neurons. The dual innervation of the esophagus collectively relays information including heat, cold, and chemicals. Vagal neurons are also known to relay vital information related to bodily homeostasis, including monitoring chemical and mechanical stimuli in the esophagus. Specifically, dorsal motor vagal neurons regulate esophageal stretch and distension, which form intraganglionic laminar endings required for swallowing (14). Disruptions in these subtypes potentially explain symptoms of esophageal dysmotility, achalasia, and dysphagia that many EoE patients experience (14).

While the innervation of the esophagus has been preliminarily described, the distinct role of esophageal afferents in allergic response and pain development remains largely unknown. Herein, we aimed to gain early insight into the neuroimmune processes that could be operational in the esophagus during type 2 immunity. In this study, we revealed elevated innervation density, neuronal sensitivity, and non-evoked pain-like behavior in allergen-challenged mice. Additionally, allergen-treatment induced marked changes in gene expression in DRGs, highlighting the impact of allergic inflammation on neuron function, particularly related to transporter genes. In an effort to expand upon these differences and identify a potential neuro-immune circuit, we explored the role of IL-4 signaling. IL-4 rapidly sensitized DRG neurons to noxious stimuli, increasing the number of neurons that respond to capsaicin and ATP. We further demonstrate that sensory neuron-intrinsic activation of this signaling pathway (*IL-4Ra*) is necessary for elevated NaV1.8+ neuronal innervation density and end-organ responses in the esophagus including the severity of allergic inflammation. These findings underscore the potential importance of a neuroimmune circuit that regulates inflammatory responses in the esophagus, having implications for esophageal disorders such as EoE.

## Results

### Allergen treatment increases pain-like behavior in mice

To establish whether allergic inflammation of the lung and esophagus has any effect on pain-like behavior in mice, we performed weekly measurements to evaluate changes in behavior over the course of repeated allergen challenges. Accordingly, we exposed mice to repeat inoculations with a common allergen *Alternaria Alternata* co-delivered to the lungs and the esophagus (**Figure 1A**) (15, 16). This experimental protocol induced esophageal remodeling, including epithelial disruption and basal cell hyperplasia, and eosinophil infiltration in the esophagus, as detected by anti-eosinophil major basic protein (MBP) staining (**Figure S1A**).

**Figure 1.**
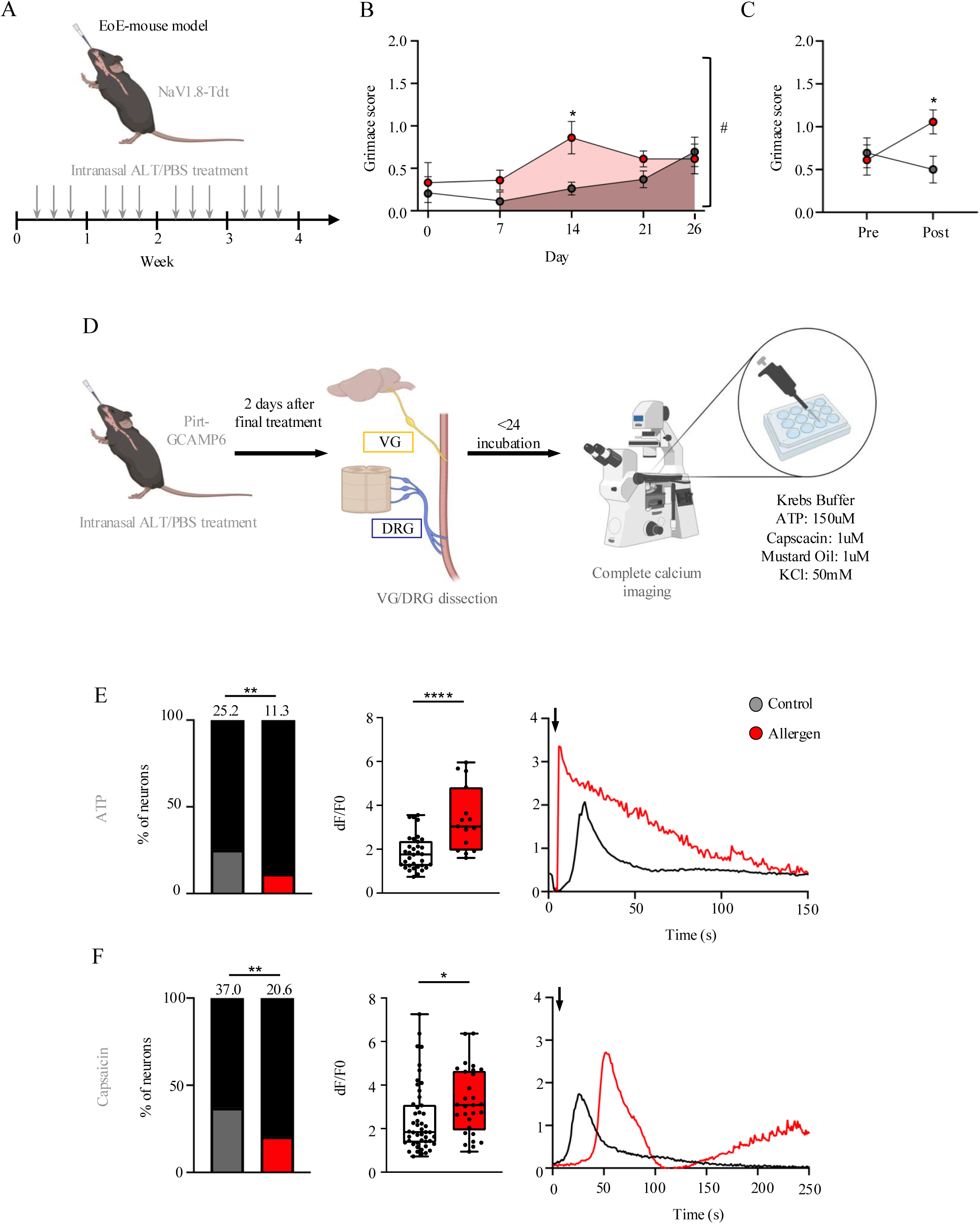
Allergen challenged mice display elevated levels of pain-like behavior and altered responsiveness of DRG neurons to noxious stimuli. (A) Schematic of the intranasal allergic esophagus inflammation model using the protease allergen *Alternaria alternata* (ALT). (B) Mouse Grimace Scale (MGS) measurements taken immediately prior to intranasal allergen challenges. Allergen treated (n=6) vs. Control (n=6). Day 14 Allergen (red) vs. Day 14 Control (gray), grimace *p<0.05 allergen vs saline at the same time point, two-way ANOVA with Holm-Sidak‘s post hoc, n=6 per group mean±SEM. #p<0.05 AUC allergen vs saline overall, student’s t-test, n=6 per group. (C) MGS measurements on day 26 immediately prior (Pre) and 20 minutes after (Post) intranasal allergen administration. *p<0.05. Two-Way ANOVA with Multiple Comparisons. Mean ± SEM. (D) Schematic of the calcium assay beginning with the allergic esophagus inflammation model outlined in Figure 1A with Pirt-GCAMP6 mice. Dissection, dissociation, and culturing of VG and T1-T4 DRG is completed two days after final ALT treatment. Imaging of cells completed <24 hours after dissection. (E-F) ATP responsive (E) and capsaicin responsive (F) DRG neurons shown in red as a percentage of total KCl-responsive neurons shown in black (left), fluorescence intensity changes (middle), and representative calcium transients from sensory neurons of PBS and ALT treated mice in response to capsaicin (1 um) or ATP (150 um) stimulation. The arrow indicates when the stimulus was added. Each dot represents one cell. (right) *p<0.05, **p<0.01, ****p<0.0001, percent responders: chi squared test, fluorescent intensity changes: unpaired T-test; DRG n>25 from ≥3 Pirt-GCAMP6 mice. Data shown as mean +/- SEM.

Allergen-treated animals displayed elevation in non-evoked pain as measured by the Mouse Grimace Scale (MGS) (17) over the course of 26 days (**Figure 1B)**. Mechanical hypersensitivity of the epigastric region was also tested following application of von Frey probes to the area (1g and 4g). In contrast to MGS scoring, there was no change in withdrawal responses to mechanical stimulation (**Figure S1B-C)**. On day 26, behavioral assays were completed before and after the final intranasal treatment to investigate the immediate impact of allergen and saline administration. Allergen-treated mice had a significantly higher MGS score compared with saline controls only after allergen (**Figure 1C**). This suggests a tight connection between the non-evoked visceral pain response to the time of allergen exposure.

### Neuronal responsiveness to noxious stimuli is modified in allergic mice

Immune cells can modulate sensitivity in neurons and our behavioral data suggests an impact of allergic inflammation on neuronal firing (2, 18–20). As such, we hypothesized that esophageal allergic inflammation may increase sensitivity of neurons to noxious stimuli. Recent single-cell analyses shows that the majority of sensory neurons from the DRG and VG express TRPV1 and TRPA1, with lower expression of P2X receptors (21, 22). As such, we examined whether TRPV1, TRPA1, and P2X expression changed under allergic conditions. Immunohistochemical (IHC) staining revealed widespread expression of TRPV1 and TRPA1, and lower expression of P2X3, but no significant differences were seen between treatment conditions (**Figure S2A-B)**. We next investigated whether VG and DRG sensitivity to capsaicin, mustard oil, and ATP was altered in allergen-treated mice. These stimuli activate the TRPV1 channel, TRPA1 channel, and P2X receptors, respectively (23–25). To test this possibility, *in vitro* calcium imaging was used on the DRG and VG from mice containing a genetically encoded calcium indicator (GCaMP6f) expressed in sensory neurons (PirtCre;LSL-GCaMP6) (**Figure 1D)**. Here, GCaMP6 is specifically expressed in >95% of all DRG/VG neurons allowing live imaging intracellular Ca^2+^ measurements as an indirect measure of action potential firing. Compared with control mice, allergen-challenge led to a decrease in the percentage of DRG cells responding to ATP and capsaicin (ATP: 25.2% vs 11.3% p<0.01; cap: 37.0% vs 20.6% p<0.01) (**Figure 1E-F**); in addition, isolated DRGs from allergen-challenged mice demonstrated increased calcium responses to both stimuli, as measured by the difference in peak (F) and baseline (F0) fluorescence (dF/F0) (ATP: p<0.0001; cap: p<0.05) **(Figure 1E-F).** This contrasts with VG neurons, where there was only a significant increase in the percentage of cells responding to ATP, and decrease in calcium response to capsaicin (**Figure S2C-D).** Neither DRG nor VG showed a difference in their response to mustard oil under allergic versus control conditions **(Figure S2E-F)**. Together, these results demonstrate that neuronal sensitivity and responsiveness to noxious stimuli in murine nerve cells is impacted by allergen-treatment.

### Dorsal root ganglia display changes in their transcriptome in response to allergen exposure

Due to the changes in neuron response seen after allergen exposure, we analyzed transcriptional changes by bulk RNA sequencing on whole DRG and VG, as well as the corresponding esophagus, under homeostatic and inflammatory conditions. We hypothesized that distinct pathways would be regulated in the ganglia of allergen-treated mice compared with saline controls, contributing to the mechanism of allergic disease development.

Principal component analysis (PCA) showed separation primarily by the tissue type; therefore, we first compared transcriptional profiles of control VG and DRG (**Figure 2A**). Both ganglia shared approximately 80/100 of their most highly expressed genes based on average expression levels. While the DRG and VG share many genes, the two groups have over 1000 differentially expressed genes (≥1.5-fold change; P<0.05; mean TPM≥1) (**Figure 2B).** This finding is in line with the previous reports that vagal neurons are largely different from the somatosensory neurons of the DRG which contributes to their unique functions (21, 26).

**Figure 2.**
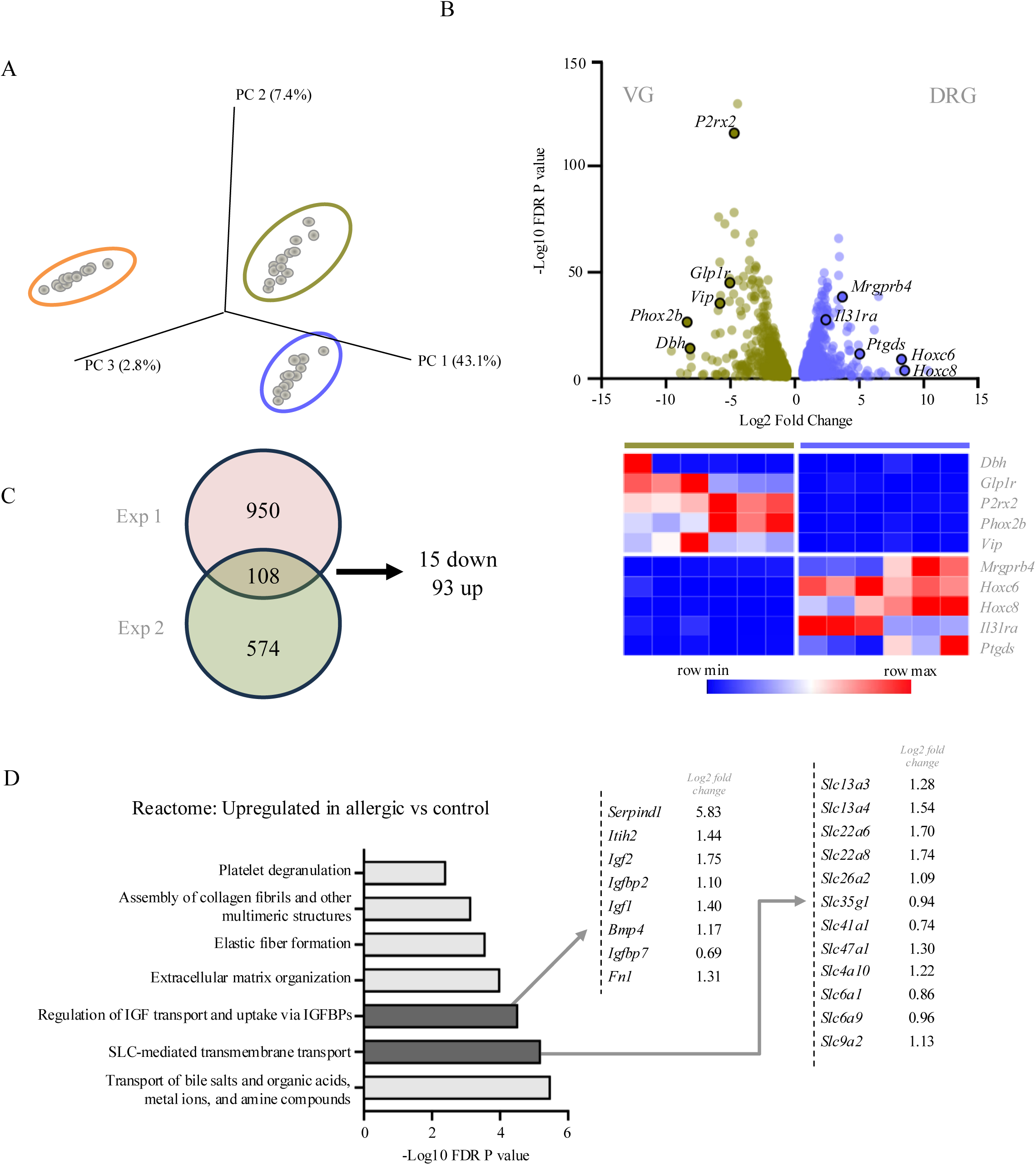
Whole genome transcriptomic analysis reveals differentially regulated genes in DRG of allergic mice. (A) Esophagus (*orange*), DRG (*green*) and VG (*purple*) transcriptome data reduced to a 3-dimensional presentation by multidimensional scaling analysis for visual presentation of the expression distance between tissue types. (B) Top: Volcano plot of the 1572 genes differentially expressed between the VG (*green*, *left)* and DRG (*purple, right)* cells. For differentially expressed genes (DEGs), FC threshold of 1.5 and p-value <0.05 between two experiments was applied. Bottom: Heatmap of ten differentially expressed genes between VG (*left)* and DRG (*right)* controls. Each column represents one mouse. (C) Venn-diagram demonstrating the number of differentially expressed genes in allergen-treated versus saline control mice in two independent experiments (exp 1: top; exp 2: bottom). DEGs that fit threshold criteria in both experiments are shown in the center; 15 downregulated and 93 upregulated. (D) Shown are the 7 most significant terms identified for the 93 upregulated genes in allergen treated DRG compared with saline controls by functional Reactome analysis. Genes for the selected terms (*grey bars*) are listed with the corresponding fold change in expression for allergen-treated mice compared to control samples. The x-axes represent the negative log (10) p-value.

While no significant difference was found between VG in the untreated and allergen treated mice, unsupervised clustering analysis on the DRG showed separation between the allergen treated and the saline control DRG (**Figure S3A)**. Two independent experiments revealed overlap of 108 differentially expressed genes between the two conditions (≥1.5-fold change; P<0.05; TPM≥1) (**Figure 2C**). Of these genes, 93 were upregulated and 15 were downregulated in the allergen versus the saline treated mice. A total of 19 genes were modified over 5-fold in at least one experiment, including genes known to be involved in allergic inflammation, *H2Q1, Serpind1,* and *Ptgds*. We aimed to identify the functional pathways regulated by allergic inflammation in DRGs. Reactome, a functional enrichment analysis database (27), revealed that the upregulated genes were enriched in multiple transport mechanisms including transport of bile salts and organic acids (FDR adjusted P = 3.22E-6) as well as Solute Carrier (SLC)-mediated transmembrane transport (FDR adjusted P = 6.25E-6), containing 12 upregulated *Slc* genes (**Figure 2E)**. Additionally, insulin-like growth factor (IGF) related genes including *Igf1, Igf2, and Igfbp2* were increased (**Figure 2E).** Notably, despite the development of inflammation in the lung and esophagus of these mice, the isolated DRGs did not demonstrate molecular changes in genes classically involved in type 2 immunity. These data demonstrate that allergen challenge in peripheral organs evokes molecular and transcriptional changes in the DRG.

### Myelinated and unmyelinated fibers differentially innervate distinct layers of the murine esophagus

We then aimed to better understand the distribution of sensory nerves in the esophagus with the ultimate goal of understanding how the distribution was affected by allergic inflammation. Whole-mount immunofluorescent analysis of murine esophagi enabled visualization of myelinated nerves via immunostaining for the neuronal axon marker BIII-tubulin. Additionally, putative nociceptive sensory axons were examined by tracking the expression of the tdTomato reporter gene under the control of the NaV1.8 promoter (28). NaV1.8+ fibers were found to be densely packed in the lamina propria, especially in juxtaposition with the epithelium, with around 6% of the total nerve signal less than 200nM from the epithelial folds (**Figure 3A**). In contrast, BIII-tubulin+ nerves were widely dispersed in the outer segments of the esophagus, significantly distal from the epithelium. This is evident in the cross-sectional views shown in **Figure 3B and 3C**. Comparing the distribution of neuronal subtypes quantitatively, NaV1.8+ fibers had a greater volume of signal closer to the basal epithelium relative to BIII-tubulin+ fibers (Median 73.52 mm (NaV1.8) vs 85.65 mm (BIII); p<0.0.1) (**Figure 3C).** NaV1.8+ fibers tracked along the length of the esophagus in a linear fashion (**Figure 3A**), whereas BIII-tubulin+ fibers formed a web-like structure around the outside (**Figure 3D, Video S4A**). In addition, localized to the layers between the muscularis and lamina propria, both NaV1.8 and BIII-tubulin nerves follow interconnecting strands of enteric neurons, forming the myenteric plexus (**Figure S4B-C)** (29). Together, this detailed mapping of murine esophageal innervation, demonstrating dense sensory nerve infiltration, prompted us to explore for their role on the impact of allergic responses.

**Figure 3.**
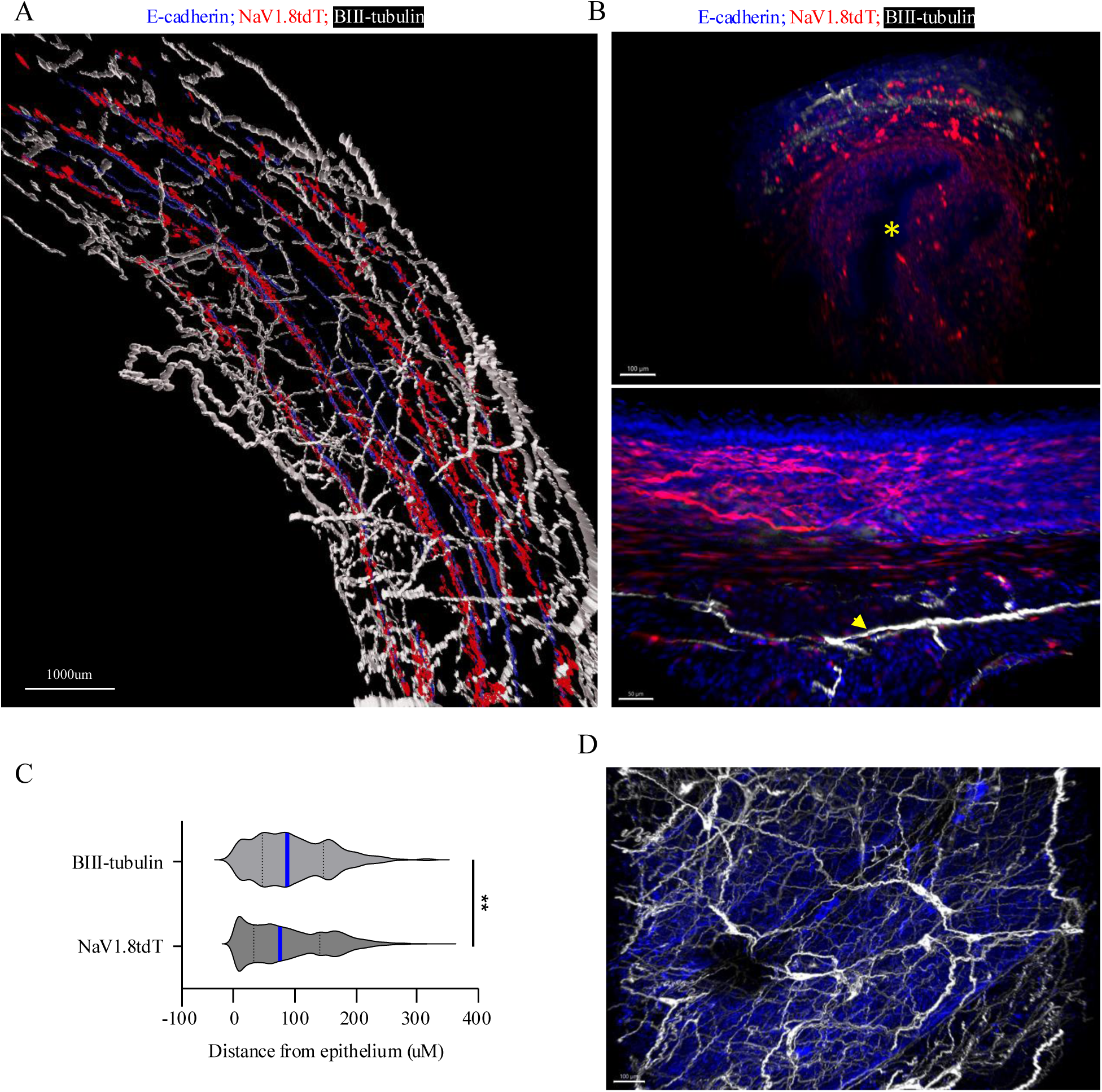
Immunofluorescent images reveal distribution of esophageal innervation in whole mount murine esophagus. (A) Whole esophagus of BIII-tubulin in NaV1.8-Cre R26CAG-floxStop-tdTomato mice. Shown are surfaces of NaV1.8+ nerves (red) within 200nm of basal epithelium (blue) with BIII-tubulin (white) around the perimeter. (B) (*top*) Cross-sectional slice of whole esophagus from NaV1.8 R26CAG-floxStop-tdTomato co-stained with BIII-tubulin (white). Yellow asterisk labels the lumen of the esophagus. (*bottom*) NaV1.8+ and BIII-tubulin+ signal in whole murine esophagus section demonstrating the distinct distribution of nociceptive (red) and myelinated (white) nerves. (C) Distribution of BIII-tubulin and NaV1.8tdT nerves as measured from the basal layer of the epithelium. Blue line designates the median. **p<0.005, Mann-Whittney test. (D) Immunofluorescence image of BIII-tubulin (white) in murine whole mount esophagus.

### Allergen treatment increases sensory neuron innervation density in the murine esophagus

After defining the esophageal innervation under homeostatic conditions, we examined the consequences of allergic inflammation on innervation. Sequential longitudinal sections allowed quantification of nerves in individual esophageal layers (**Figure 4A, Figure S4D)**. Allergen-treated mice exhibited increased density of NaV1.8-Tdtomato nerves in the lamina propria (1.7-fold increase, p<0.0001) compared with controls (**Figure 4B)**. In contrast, there was not a significant difference in myelinated fiber density in any esophageal layer between the two groups (**Figure 4c).** Taken together, these findings indicate that allergen-induced esophagitis increases nociceptor innervation density specifically in the lamina propria of the murine esophagus.

**Figure 4.**
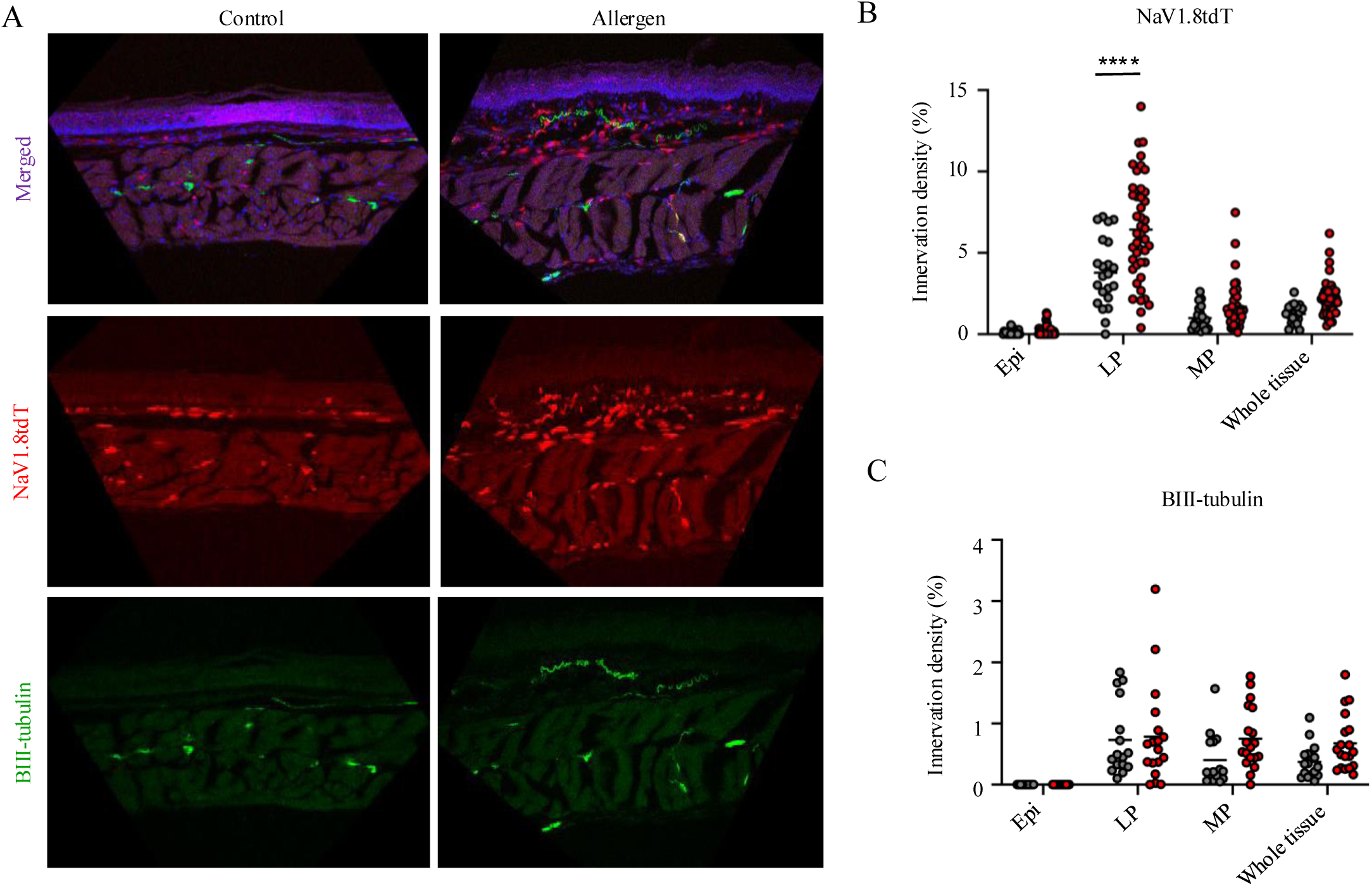
Increased esophageal sensory neuron innervation density in the lamina propria of allergen-treated mice. (A) Representative immunofluorescence images of BIII-tubulin in longitudinal esophageal cuts from NaV1.8-Cre R26CAG-floxStop-tdTomato mice treated with PBS and intranasal ALT. (B) Murine Esophageal NaV1.8-tdT+ nerve area; LP tissue control vs Alternaria treated, ****p<0.0001. Data were obtained from 3 independent experiments; allergen treated (n=8) vs. Control (n=4). (C) Murine Esophageal BIII-tubulin nerve density. Data were obtained from 2 independent experiments; allergen treated (n=6) vs. Control (n=5). Individual data points represent a separate section of esophagus. Two-way ANOVA with multiple comparisons, horizontal line shows mean.

### Patients with active EoE have increased nerve density

Next, we aimed to determine if changes in innervation pattern might occur in patients with EoE. Whole-mount immunofluorescent staining of the neuronal axon marker BIII-tubulin in the human esophagus enabled visualization of myelinated nerves revealing areas of dense innervation most notably in the lamina propria near the basal epithelium (red arrowheads, **Figure 5A**). While these fibers have increased density near the lumen, BIII-tubulin+ fibers were distributed throughout the entire width of the esophagus (**Figure 5A and 5B).** Whereas murine esophagi showed little BIII-tubulin fluorescent signal in the lamina propria, the staining was in proximity to the basal epithelium in humans (**Figure 5B**). Three-dimensional imaging of the human esophagus revealed evidence of the enteric plexuses, with the submucosal plexus (yellow bracket **Figure 5A**) localized closer to the lumen, and the myenteric plexus (orange arrows **Figure 5A**) between the muscle layers of the outer esophagus. In order to investigate individual nerves and map their specific location, we shifted to analyzing consecutive sections from esophageal biopsies. Analysis of biopsies from healthy controls was consistent with whole-mount visualization, with BIII-tubulin localized mainly to the lamina propria (**Figure S5C)**. Myelinated fibers could also be traced into the papilla (**Figure 5C)** and occasionally laced in the epithelium (**Figure 5D**). Innervation density was analyzed in esophageal biopsy samples isolated from a cohort of subjects with active EoE (n=39) and healthy controls (n=11) (**Table 1)** (**Figure 5E)**. Subjects with active disease had significantly higher myelinated innervation density compared with healthy controls (**Figure 5F).** Esophageal innervation density was stratified by sex, history of atopy, and age, and remained elevated in active EoE patients compared with controls. There was no correlation between eosinophil esophageal levels and myelinated nerve density in patients with active EoE **(Figure S5D)**.

**Figure 5.**
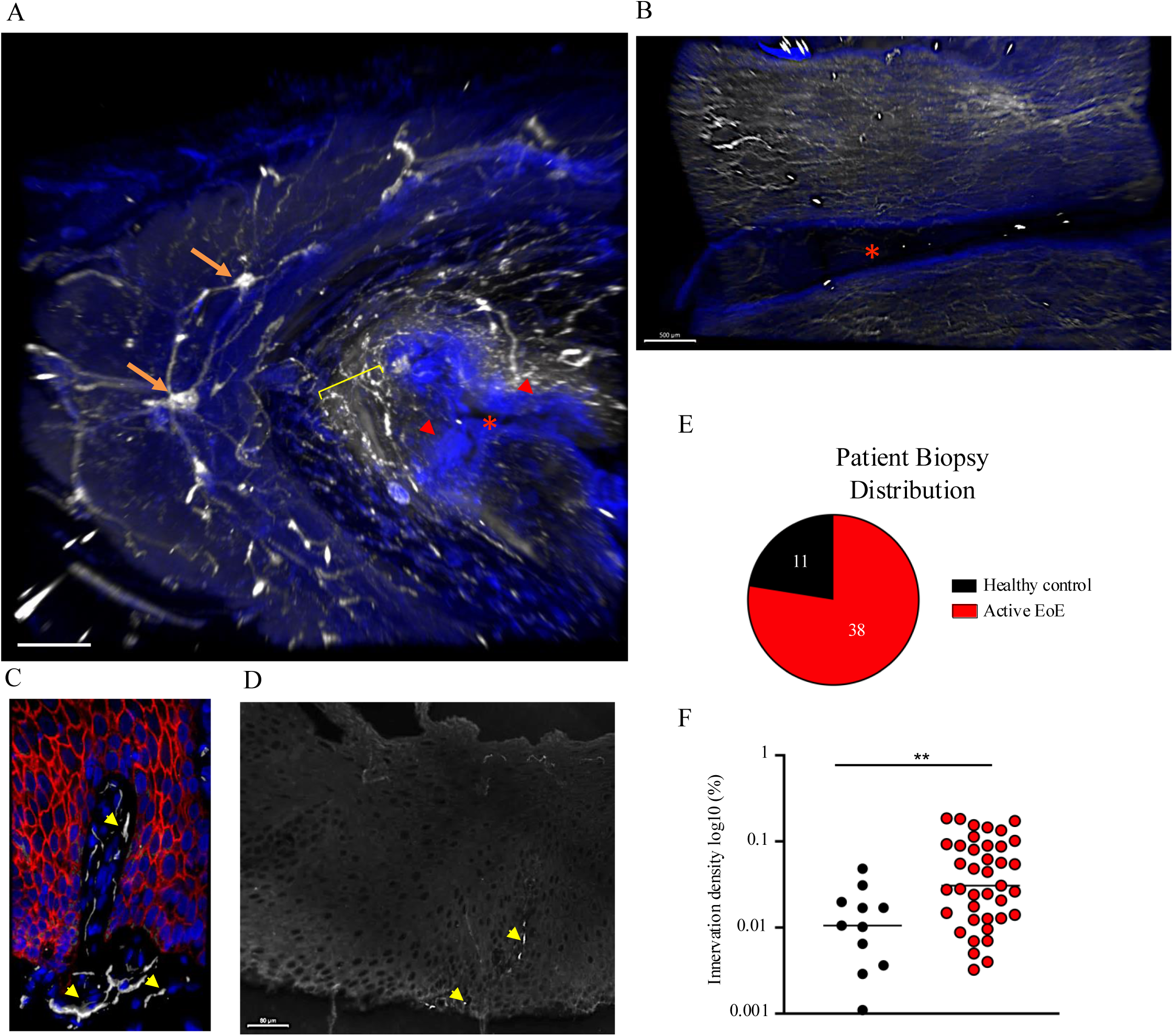
Active EoE patients have elevated myelinated innervation density in the esophagus. (A) Whole esophagus from 36-year-old donor labeled for BIII-tubulin (white) and DAPI (blue). Visualization of the myenteric plexus (orange arrows), basal epithelium (red arrowheads), lamina propria (yellow bracket), and esophageal lumen (red asterisk). (B) Slice of whole mount human esophagus labeled for BIII-tubulin (white) demonstrating the distribution of myelinated nerves throughout the entire thickness of the organ. Lumen is designated by the red asterisk. (C) Immunofluorescence image of DAPI (blue), E-cadherin (red), and BIII-tubulin (white) in esophageal biopsy from an active EoE patient, with nerves (yellow arrowheads) travelling within the papilla of the lamina propria. (D) Immunofluorescence image BIII-tubulin (white) in esophageal biopsy from an active EoE patient demonstrating evidence of nerves (yellow arrowheads) located in the epithelium. (E) Distribution of 58 patient biopsies based on current diagnosis. (F) Active EoE patients (n=38) show elevated BIII-tubulin density in their esophageal biopsies when compared with healthy controls (n=11). Each dot represents the average density from >1 esophageal section from one patient. **p<0.005, Mann-Whitney test. Data is shown as mean.

**Table 1:** Demographic data of patient cohort.

| Characteristic | Control | Active EoE | EoE Remission |
| --- | --- | --- | --- |
| No. of subjects | 11 | 38 | 9 |
| Age at biopsy (years) | 11 ± 4.4 | 11.2 ± 4.6 | 9.8 ± 5 |
| Sex (male) | 2 (18.2) | 30 (78.9) | 8 (88.9) |
| Race (white) | 9 (81.8) | 84 (89.5) | 9 (100) |
| History of atopy | 4 (36.4) | 31 (81.6) | 5 (55.6) |
| Tissue eosinophil count | 0 | 54 (15-200) | 0 |

### Neuroimmune interactions occur in the esophagus of humans and mice

Previous studies have shown the potential for both mast cells and eosinophils to interact with nerves under a variety of conditions, which may explain changes to esophageal nerves during disease (9, 29, 30). High resolution IF imaging of esophageal biopsies from EoE patients revealed direct interactions between BIII-tubulin+ nerves and mast cells, visualized with anti-tryptase antibody (**Figure 6A),** consistent with a recent short communication from Zhang et al (32). Additionally, staining for eosinophil peroxidase (EPX) showcased myelinated nerves in close proximity to EPX granules (**Figure 6B)**. Together, these imaging analyses suggest an interaction between nerves and mast cells as well as eosinophils in active EoE.

**Figure 6.**
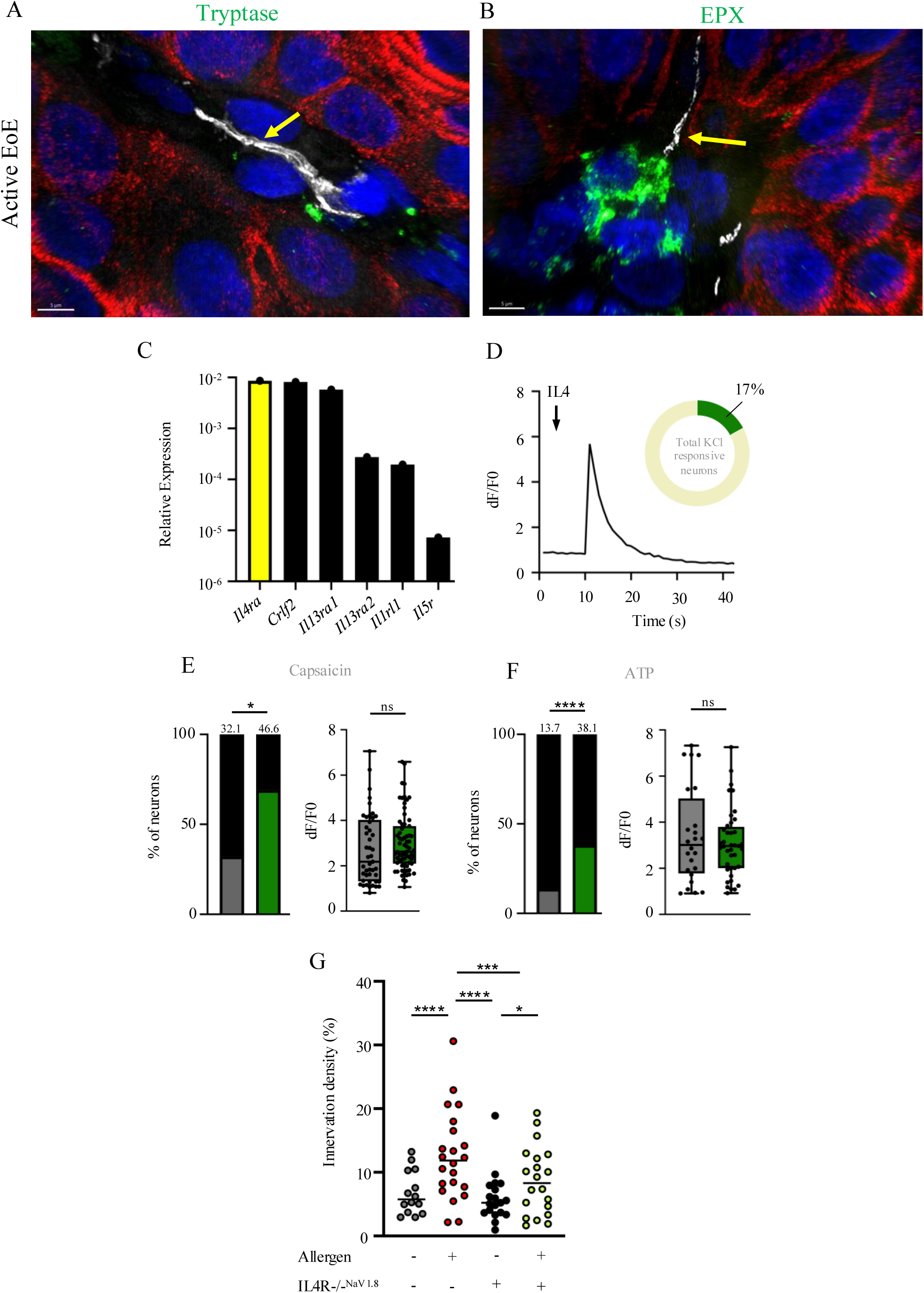
Potential neuroimmune interactions in the allergic esophagus involving IL4Ra signaling pathway. (A-B) 60x water Sora images showing interaction between BIII-tubulin+ nerves (white) and (A) tryptase (green) and (B) eosinophil peroxidase (EPX) (green) in active EoE patient esophageal biopsy. (C) Comparison of IL4Ra, CRLF2, IL13Ra1, IL13Ra2, IL1RL1, and IL5R, expression by qRT-PCR in whole mouse naive DRG n = 4 WT mice. (D) Representative calcium transient of whole DRG calcium response to recombinant mouse IL-4 (300nm). Graph outlines total number of KCl responsive cells to IL-4 (67/392). (E-F) (E) Capsaicin responsive and (F) ATP responsive DRG neurons as a percentage of total KCl-responsive neurons (left) and fluorescence intensity changes (right), in response to capsaicin (1 um) or ATP (150 um) stimulation after vehicle (gray) or IL-4 (300nm) (green) exposure. Each dot represents one cell. *p<0.05, ****p<0.0001, percent responders: chi squared test, fluorescent intensity changes: unpaired T-test; DRG n>100 from 3 Pirt-GCAMP6 mice. Data shown as mean +/-SEM. (G) Murine esophageal Nav1.8tdT nerve density. Data were obtained from 2 independent experiments; WT control, WT allergen treated, IL-4R-/-^NaV1.8^ control, and IL-4R-/-^NaV1.8^ allergen treated. n≥4 mice per group; individual data points represent a separate section of esophagus. Two-way ANOVA with multiple comparisons, horizontal line shows mean.

Various nociceptors have been shown to respond to cytokines known to be produced by mast cells and eosinophils, including cutaneous and muscle nociceptors to IL-1β (32, 33), and lung nociceptors to IL-5 (35). More specifically, type 2 cytokines can also modulate the responses of neurons to subsequent exposure to pruritogens (6, 8). Therefore, we hypothesized that similar signaling could occur in our experimental EoE model and explain the observed functional, transcriptomic, and innervation changes. To explore potential candidates for this neuroimmune communication pathway, we examined the transcript profiling of naïve DRG and found expression of cytokine receptors, consistent with recent data (8). This included *Il-4ra, Crlf2, Il-13ra1, and Il-5r* (**Figure 6C**). *Il-4ra* had the highest expression levels on naïve DRG neurons compared with other cytokine receptors. This receptor and its signaling pathway are of high interest, as EoE pathogenesis has been shown to be dependent on overexpression of the IL4Rα and its ligand IL-13; anti-IL-4Ra is now an approved therapy for EoE (35, 36). Together, these results highlight the potential for neuroimmune interactions in EoE, and provide a variety of starting points for probing the mechanism of the changes that we have demonstrated so far.

### IL-4 alters DRG responses to capsaicin and ATP

Signaling via IL-4Rα has also been shown to directly activate sensory neurons through TRP channel-dependent calcium influx (38), potentially explaining the changes in capsaicin responses that we observed (**Figure 2F**). Therefore, we hypothesized that similar signaling could occur in experimental EoE and directly alter DRG responsiveness to capsaicin and ATP. We explored IL-4R activation in cultured DRG neurons and tested if IL-4 exposure had any effect on neuronal response to capsaicin and ATP. Exposing naïve DRG neurons to IL-4 (300nM) produced calcium responses in 17% of neurons (67/392 KCl+ cells, 17.09%), matching previous reports **(Figure 6D)** (6). IL-4 exposure was shortly followed by exposure to low concentrations of capsaicin or ATP; responses were measured and compared to vehicle exposed DRGs. IL-4 sensitized neurons demonstrated increased numbers of cells responding to both capsaicin (46.6% vs 32.1%, p<0.05) and ATP (38.1% vs 13.7%, p<0.001) (**Figure 6E-F).** No differences in peak calcium were seen in response to capsaicin or ATP. We also incubated the DRG neurons in media with IL-4 overnight and repeated calcium imaging 24 hours later. Prolonged exposure to IL-4 led to a reverse in the sensitization seen with acute IL-4 exposure (**Figure S6A-B).** Our findings demonstrate the ability for IL-4 to immediately sensitize DRG neurons to capsaicin and ATP, but after prolonged exposure this effect is diminished. This highlights the potential for IL-4 to directly contribute to acute hypersensitivity in neurons.

### Sensory neuron IL-4Rα contributes to elevated innervation density in allergen treated mice

IL-4 has be shown to facilitate nerve regeneration and regrowth (38, 39). As such, we hypothesized that IL4 receptor ablation on sensory neurons would reduce the nociceptive innervation density increases seen in the allergen treated mice. To test this hypothesis, we generated mice lacking IL4Rα specifically on NaV1.8 neurons (NaV1.8-Cre^+^ IL-4R^fl/fl^ tdTomato; IL-4R-/-^NaV1.8^). Following allergen or control saline exposure, quantification of NaV1.8 nerve density in WT and IL-4R-/-^NaV1.8^ mice revealed innervation differences (**Figure 6G)**. Allergen challenged WT mice demonstrated elevated innervation density in their lamina propria compared with WT (p<0.0001) and IL-4R-/-^NaV1.8^ saline controls (p<0.0001). Notably, IL-4R-/-^NaV1.8^ allergen-challenged mice displayed significantly less nerve density compared with WT allergen-challenged mice (p<0.001) and saline treated control IL-4R-/-^NaV1.8^ mice (p<0.05). Together, these findings demonstrate that neuronal IL-4Rα signaling regulates allergen-induced increases in sensory neuron innervation density.

### Disruption of neuronal IL-4Rα signaling alters esophageal inflammatory environment and whole DRG transcriptome in allergic mice

Next, we hypothesized that disrupting IL-4Rα neuroimmune signaling might decrease type 2 inflammation in the esophagus of allergen challenged mice. Accordingly, we examined bulk-RNA sequencing of the esophagus from IL4R*-*/-^NaV1.8^ and WT mice, both including mice with and without allergen challenges. Initial comparison between WT allergic and saline control mice revealed 582 differentially expressed genes between the two conditions (≥1.5-fold change; P<0.05; TPM≥1) (**Figure 7A**). Of these genes, 481 were upregulated and 101 were downregulated in the allergen versus the saline treated mice, including genes selective to eosinophils (*Rnase2a* and *Ccr3*), and involved in the innate immune response (*Chil3, Chil4, Ccl8*) (**Figure S7A**). Reactome pathway analysis of the RNA sequencing (RNA-seq) data showed that expression of immune system- and inflammation-associated pathways was significantly increased in WT allergic mice compared with saline control WT mice (**Figure 7B**). Allergen-challenged IL-4R-/-^NaV1.8^ mice exhibited differential expression of 61 genes compared with allergen-challenged WT mice (≥2-fold change; P<0.05; TPM≥1). A total of 31 genes were also modified between the WT saline control and allergic mice, suggesting a role for IL-4R-/-^NaV1.8^ signaling in expression level of this subset of genes (**Figure S7B**). STRING biological processes analysis of the 25 upregulated genes revealed the expression of defense and immune response pathways were significantly upregulated in allergic WT mice compared with allergic IL-4R-/-^NaV1.8^ (**Figure 7C).** Differentially expressed genes included those related to eosinophil chemotaxis (*Ccr3*), eosinophil-derived cytotoxicity (*Rnase2*), and allergic inflammation (*Alox15*); select genes were validated with an additional cohort via qPCR (**Figure S7C**). The pathological contribution of neuronal IL-4Rα was examined by detailing allergen-induced histopathology in the esophagus. Allergen-challenged IL-4R-/-^NaV1.8^ mice exhibited markedly reduced esophageal pathology as evidenced by decreased eosinophil infiltrates, basal cell hyperplasia and an intact epithelium compared with allergen-challenged WT mice (**Figure 7D and S7D-E)**.

**Figure 7.**
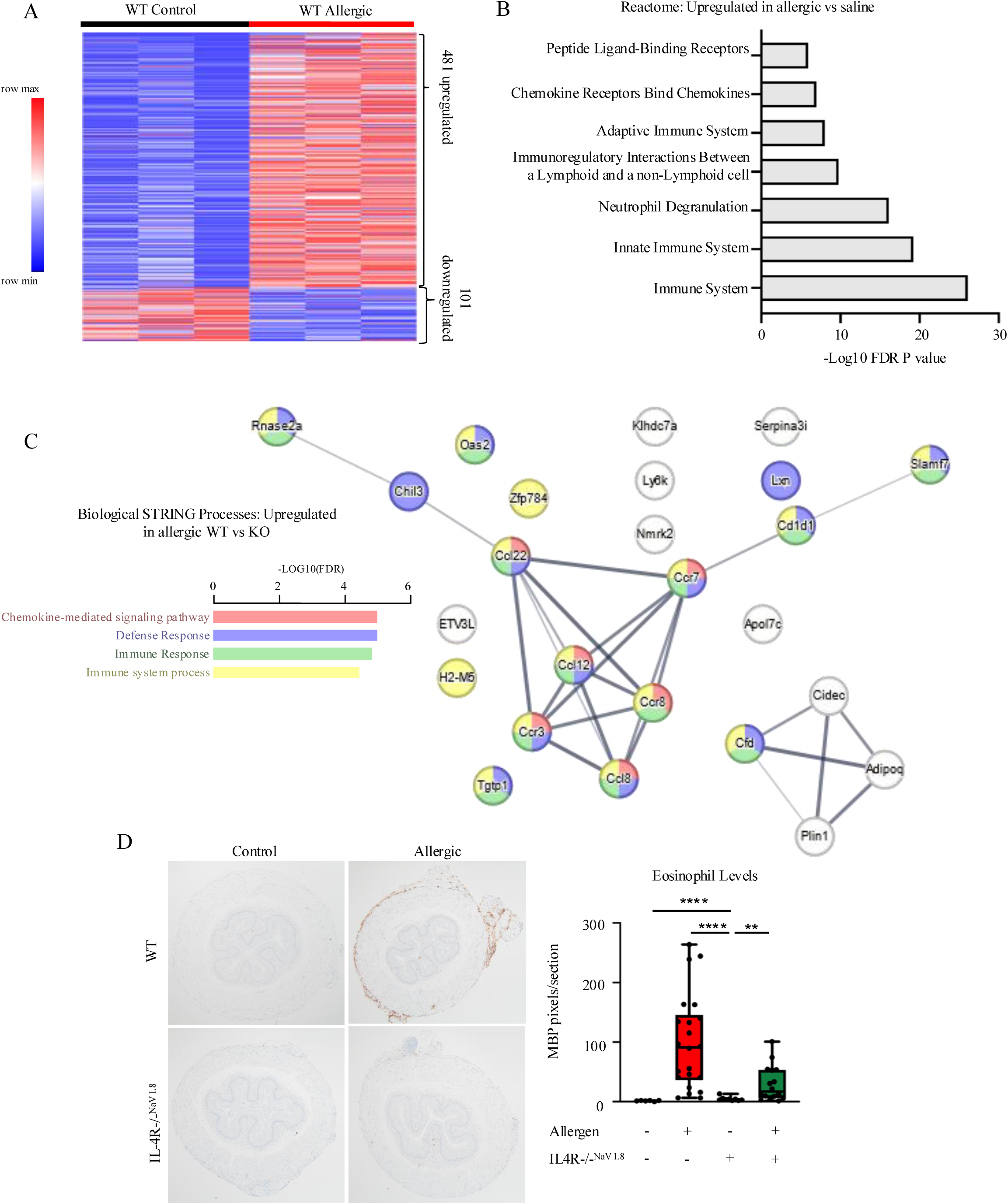
Neuronal IL-4Ra Expression Is Necessary for Esophageal Eosinophilic Inflammation. (A) Heat map based on 582 differentially expressed genes in control vs allergen challenge esophagus. Upregulated genes (red) vs downregulated (blue) of expression profiles of differentially dysregulated genes. ≥1.5-fold change; P<0.05; TPM≥1; Each column represents one mouse (B) Reactome analysis of 481 upregulated genes in the esophagus of allergen treated mice. Data shows top 7 pathways based on FDR p-value. (C) Biological STRING process analysis based on 25 upregulated genes in allergen challenged WT vs IL-4R-/-^NaV1.8^ mice. Colors of circles correspond with pathways of the same color. Line thickness indicates the strength of data support. (D) Representative immunohistochemistry staining of eosinophil major basic protein (MBP) staining from WT control, WT allergen treated, IL-4R-/-^NaV1.8^ control, and IL-4R-/-^NaV1.8^ allergen treated. (E) Eosinophil major basic protein (MBP) pixel quantification per esophageal section comparing WT control, WT allergen treated, IL-4R-/-^NaV1.8^ control, and IL-4R-/-^NaV1.8^ allergen treated mice. n≥2 mice per group; individual data points represent a separate section of esophagus. One-way ANOVA with multiple comparisons, horizontal line shows mean.

Finally, we aimed to examine what signals originating from sensory neurons may be altered due to the presence or absence of IL4Rα, potentially explaining the changes in esophageal inflammation and sensitivity that occurs during allergen exposure. Bulk-RNA sequencing of DRGs from in IL-4R-/-^NaV1.8^ and WT allergen-challenged mice revealed 224 differentially expressed genes between the two groups (≥1.5-fold change; FDR<0.05; TPM≥1). Of the 224 genes, 215 were downregulated in the IL-4R-/-^NaV1.8^ allergic vs WT allergic DRG (**Figure 8A**). Reactome analysis on the 215 downregulated genes revealed a decrease in the formation of multiple structural components including collagen elastic fiber formation, integrin cell surface reactions, and extracellular matrix organization (**Figure 8B**). Many of these pathways were also found to be enriched in the WT allergic DRG when compared to saline controls (Figure 2D). Additionally, of the 108 differentially expressed genes identified between WT allergic and saline DRGs (Figure 2), 56 were also different between IL-4R-/-^NaV1.8^ and WT allergic DRGs (**Figure S8A-C**). This significant overlap provides insight into the potential role of IL4Rα signaling in mediating allergic inflammation through sensory neurons. Of note, this included insulin-like growth factor (IGF) related genes including *Igf1, Igf2, and Igfbp2* (**Figure 8B).** Together, these data reveal an influential IL-4Rα neuroimmune circuit that regulates allergic inflammation in the esophagus and is potentially mediated by differential sensory neuron mediator release.

**Figure 8.**
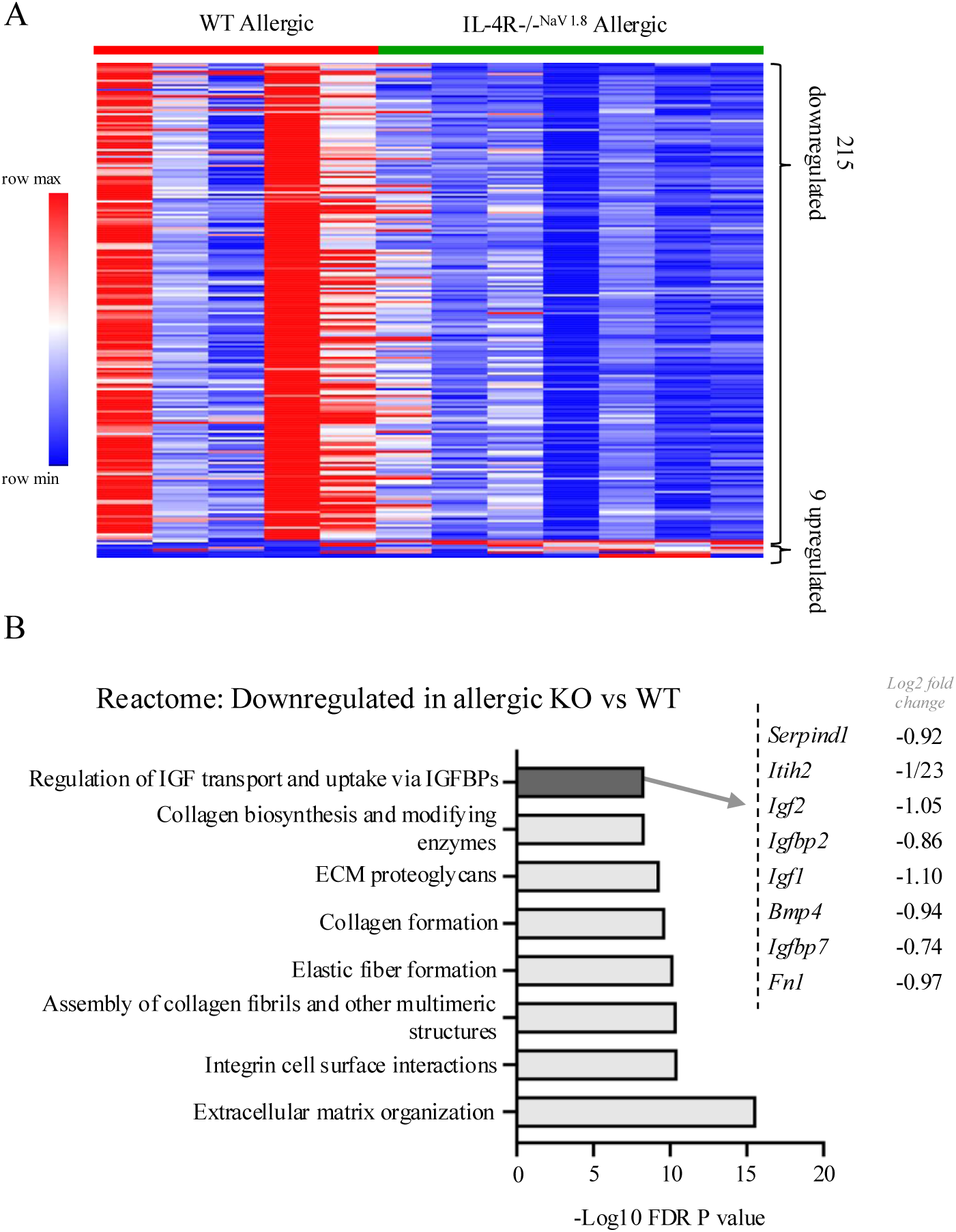
IL-4R-/-^NaV1.8^ allergen-treated mice have a distinct DRG transcriptome compared to WT allergen-treated mice. (A) Heat map based on 224 differentially expressed genes in DRGs from WT allergen treated mice vs, IL-4R-/-^NaV1.8^ allergen treated mice. Upregulated genes (red) vs downregulated (blue) of expression profiles of differentially dysregulated genes. ≥1.5-fold change; P<0.05; TPM≥1; Each column represents one mouse. (E) (*left)* Reactome analysis of 215 downregulated genes in the DRG of IL-4R-/-^NaV1.8^ allergen treated mice compared with WT allergen treated. Data shows top 8 pathways based on FDR p-value. (*right)* Genes for the selected term (*grey bar*) are listed with the corresponding fold change in expression for allergen-treated IL-4R-/-^NaV1.^ mice compared to WT allergen treated samples. The x-axes represent the negative log (10) p-value.

## Discussion

Here, we decipher communication between inflammation and the nervous system in the context of allergic responses in the esophagus via IL4Ra-signaling. EoE-like inflammation elevated non-evoked pain-behavior in mice and modified neuronal responsiveness to ATP and capsaicin, increasing the peak response in DRG cells to both stimuli (Figure 1). Not only did allergic inflammation alter neuronal physiology, whole DRG transcriptome analysis revealed differentially regulated genes in allergen exposed mice compared with saline exposed controls (Figure 2). In mice, nociceptive nerves were found to be densely packed near the epithelium, while myelinated nerves were localized to the outer segments of the esophagus (Figure 3). This innervation was found to be modified following repeated allergen challenges, with elevated nociceptive neurons located in the lamina propria (Figure 4). In addition, we identified a type 2 neuroimmune circuit that involves the interplay sensory-neuron specific IL4Rα which modifies esophageal end-organ innervation (Figure 6) and inflammatory response (Figure 7) and modifies DRG gene expression (Figure 8). Translational studies provided evidence for neuroimmune interplay in human EoE (Figure 5). Taken together, our findings provide evidence for an allergen-induced neuroimmune loop responsible, at least in part, for esophageal inflammation and proprioception.

Patients who suffer from EoE experience a variety of symptoms including overall discomfort, abdominal and chest pain, and increased pruritis (41). Previous work has evaluated the potential role for peripheral neurons in the development of symptoms and inflammation of other allergic diseases, including atopic dermatitis (AD) and allergic asthma, but little has been done to investigate this relationship in allergic esophagitis. We found DRG neurons from allergen-treated mice have fewer cells that respond to capsaicin and ATP, but the responding cells display increased peak responses to both stimuli (Figure 1). Perhaps the capsaicin and ATP responsive cells have an elevated peak calcium response in an effort to compensate for a decrease in the number of TRPV1 and P2X-cells that are being activated.

To better understand how chronic allergen-challenge impacts neuronal function at a transcriptional level, bulk RNA sequencing of whole DRG and VG was performed (Figure 2). The DRG was markedly impacted by allergen exposure, modifying the expression of over 100 genes; this is especially notable in view of the DRG not likely being directly exposed to the allergens. Functional analyses revealed changes in multiple pathways, including regulation of insulin-like growth factor (IGF) genes, including upregulation of the genes *Igf1, Igf2,* and *Igfb2*.

Of note, we previously demonstrated that IGF-1 signaling regulates neuronal sensitization and pain (42). Additionally, increased expression of IGF-1 enhances the regeneration of neurites from DRGs, increasing neuronal viability and outgrowth (43, 44). The link between elevated *Igf1* in allergic DRG, and the role of *Igf1* in neuronal growth, may be an additional factor contributing to the altered DRG response (Figure 1) and increased sensory innervation seen in mice with under the same conditions (Figure 4). Interestingly, the IGF-1 receptor, IGF1R, is upregulated during eosinophil development and receptor levels are regulated by miR-223 which is highly upregulated in patients with EoE (45), suggesting a potential relationship between DRG and eosinophils in EoE pathogenesis via the IGF-pathway. This pathway has also been shown to have an important role in allergic airway remodeling and esophageal basal-cell hyperplasia via upregulated *Igfbp3* (46, 47). Together, whole transcriptome analysis on the DRG and VG showcased specific changes occurring in allergic mice when compared with healthy controls, mainly localized to the DRG. This highlights, that while the esophagus is innervated by both ganglia, each has a distinct role. The top differentially expressed genes between the control DRG and VG call attention to their unique functions. For example, the DRG contained genes known to fall into specific sensory subtypes including *Il31ra* which is expressed by NP3 neurons. This subtype is labeled by the expression of natriuretic peptide type B (NPPB), which serves as a neurotransmitter to signal itch information to spinal cord relay neurons (22, 48). Another highly expressed gene in the DRG included *Mrgprb4,* unmyelinated, nonpeptidergic nerves, which has been shown to transmit mechanical information (49). This subtype is also predicted to be involved in G protein-coupled receptor signaling pathway, mast cell degranulation, and positive regulation of cytokinesis.

Converging data of EoE pathogenesis has placed the esophageal epithelium at the center of the disease (50). Not only is the epithelium thought to be a driver of disease progression, it is also the location of the elevated inflammatory infiltrates (52, 53) including eosinophils and mast cells during EoE (52). We provide evidence of the proximity of the immune infiltrate and sensory nerves (Figure 6), suggesting the potential for intimate communication between the two systems. This interaction likely includes crosstalk between these allergic inflammatory eosinophils and nerves. It is notable that eosinophils have been shown to increase sensory nerve branching in humans and mice under various allergic conditions, consistent with the presented findings (29, 55). One potential interacting set of mediators are the IGFs, which have been shown to promote the proliferation of immature eosinophils as well as aid in neuron migration and outgrowth (43, 45, 56). Another potential interaction could include the binding of type 2 cytokines to their corresponding receptors on the sensory neurons themselves. It has been shown that type 2 cytokines, including IL-4 as shown, can directly stimulate neurons in humans and mice (38). IL-4 has been implicated in nerve regeneration and regrowth following injury (38, 39), collectively supporting our finding that IL4Rα is necessary for elevated esophageal innervation in mice (Figure 6).

In order to expand the murine findings, we demonstrated that the esophagus of subjects with active EoE have elevated myelinated nerve density. Whole mount analysis of both human and murine esophagi revealed differing distributions of myelinated nerves throughout the organ. In mice, BIII-tubulin+ nerves were localized further from the epithelium in contrast to the nociceptive fibers (Figure 3). Contrary, myelinated fibers in the human esophagus were seen in every layer, including in the epithelium (Figure 5). The differences in nerve distribution under homeostatic conditions may explain the innervation changes in mice vs humans; as the nerves that lie closer to the epithelium look to be more highly impacted. It is notable that the murine and human esophagus have unique epithelium, the former being keratinized, which may have impact on the need for various nerve innervations.

Frequent dosing (weekly) of anti-IL-4Rα monoclonal antibody therapy improves both EoE histologic outcomes and symptoms but less frequent dosing (biweekly) only improves the histopathology (55). Our findings raise the possibility that IL4Rα signaling on sensory nerves may be a contributing factor to EoE, and elicitation of different IL-4Rα responses may require variable levels of inhibition. IL4Rα has been previously evaluated in the setting of atopic dermatitis, where neuronal IL-4Ra signaling was found to critically mediate chronic itch (38). Without the receptor on sensory nerves, scratching, ear thickness, and histology grade are diminished suggesting a pro-inflammatory role for neuronal IL4Rα. Kim also et al. revealed that type 2 cytokines directly activate murine sensory neurons, and IL-4 can increase responses in subsets of neurons to a variety of pruritogens (38). Deciphering the role of this receptor may provide insight into the results seen in the clinical trials and provide building blocks for additional treatment strategies, yet the effect of IL4Rα neuroimmune signaling on esophageal allergic inflammation remains unknown.

Consistent with published data, the calcium imaging studies indicate that neuronal responses to IL-4Rα stimulation occur within seconds to minutes, sensitizing DRG-neurons almost immediately to capsaicin and ATP (Figure 6). We reasoned that acute exposure to IL-4 may not replicate the *in vivo* environment accurately, as DRGs from our allergen-challenged mice showed different responses. In an attempt to recapitulate an inflammatory environment, we incubated the neurons in IL-4 overnight, but there were no changes to neuron responsiveness or peak calcium levels (Figure S6). This highlights the potential for IL-4 to directly contribute to acute hypersensitivity in neurons, but more exploration is needed to determine what else is contributing to the calcium responses seen in allergen-treated mice. Future studies will allow for a better understanding of both rapid sensitization and transcriptional changes induced by type 2 cytokines, in conjunction with other inflammatory mediators. While IL4Rα activation may not fully explain the neuronal sensitivity changes, our study demonstrates that sensory neuron IL4Rα mediates innervation density (Figure 6G) and the esophageal immune response (Figure 7). Based on the decrease of esophageal eosinophil specific gene levels and eosinophil levels via MBP staining in allergic mice without neuronal IL-4Rα, we speculate that the improvement of both histologic outcomes and symptoms in patients receiving dupilimab may be due to the inhibition of signals originating from sensory neurons. Based on our bulk-RNA sequencing analyses on DRGs, we hypothesize that changes in IGF-signaling during allergic inflammation may play a key role in mediating DRG-dependent esophageal responses. As mentioned, elevated IGF-1 may explain multiple components of the neuronal changes shown here, including elevated density (43, 44) and hypersensitivity (42). Additionally, IGF-1 has been shown to promote mast cell survival (56), regulate eosinophil levels and cause proliferation of epithelial cells in allergic asthma (57). IGF-1 release has also been tightly linked to IL4Rα activation in macrophages (58) and retinal ganglia cells (59). Thus, IGF-1 signaling may have multiple functions in allergic inflammation, dependent in part upon IL4Rα activation of sensory neurons. Future studies aimed at understanding these pathways, and other sensory neuron released signals, may reveal a IL4Rα-dependent mechanism specific to sensory neurons. Overall, a deeper understanding of how neurons respond to activation of IL4Rα, and how this subsequently impacts the inflammatory environment, may have important clinical considerations for additional treatment options adjacent to dupilimab.

There are a number of limitations to the current study that are noteworthy. The human data is limited by a small sample size with little control over the size, shape, and orientation of the esophageal biopsies. The aero-allergen model of EoE is based on artificial implementation of allergic disease and therefore may not well translate to the human disease. The comparison between mouse and human data has limitations given different tissue structure between mouse and human esophagus, such as the later not being keratinized. Additionally, the thoracic DRGs that were chosen for analysis innervate both the esophagus and the lung. Deciphering the exact physiological changes on esophageal innervating DRG neurons would need further investigation.

Understanding the bidirectional communication and interplay of peripheral nerves and inflammatory cells is a growing topic in medicine (6–9,20,60,61). Our findings provide preliminary advancement in understanding the neuroimmune system involved in allergic inflammation, especially in the context of the esophagus. We report that allergic inflammation (in mice and man) is associated with changes in esophageal nerve density as well as DRG responses as measured by ex-vivo stimulation to neuro-agonists and genome-wide molecular transcript profiles particularly related to gene encoding for transporters and response to IGFs. We highlight the potential importance for a neuroimmune circuit at work in the esophagus, connected via sensory neuron IL4Ra. These findings provide early insight into neuroimmune mechanisms that may be operational during allergic responses in the esophagus and draw attention to the potential of therapeutic strategies that target the neuro-immune axis, particularly to treat symptoms such as pain.

## Methods

### Patients

Biopsies were acquired from individuals who were having an endoscopy for EoE or related symptoms at Cincinnati Children’s Hospital Medical Center (CCHMC). Active EoE was defined by a histologic finding of 15 or more esophageal eosinophils per microscopic HPF with clinical symptoms. Controls included histologically normal individuals with no history of EoE and patients with a history of EoE with the endoscopy showing 0 eosinophils/HPF at the time of the biopsy. Individuals’ information was obtained from electronic medical records and research questionnaires. Patient data are summarized Table 1. Participants provided written informed consent for inclusion in an IRB-approved protocol. Sex was considered as a biological factor during quantification of innervation density, but did not impact the conclusions.

### Animals

Adult male and female mice with a C57BL/6 background, between 3 to 6 months of age were used throughout all experiments. All mice were randomly assigned to experimental groups in an age- and gender-matched fashion, and littermates were used whenever possible. Mice were housed in a barrier facility maintained on a 14:10-hour light–dark cycle with a temperature-controlled environment and given food/water ad libitum.

Nociceptive neuron reporter animals were generated by crossing the NaV1.8-Cre positive animal (Jax Stock#: 036564) with the tdTomato (tdTom) reporter mouse (B6.Cg-Gt(ROSA)26Sortm14(CAG-tdTomato)Hze/J) purchased from The Jackson Laboratory (Stock#: 007914). Nociceptive neuron specific IL4Ra knockout animals were generated by crossing the NaV1.8-Cre positive animal to the IL4Rafl/fl animal that was kindly donated to us by Dr. Brian Kim at Mount Sinai. Pirt-GCAMP6 animals were generated by crossing Pirt-Cre animal (kindly donated to us by Xinzhong Dong at Johns Hopkins University) to LSL-GCaMP6fl/fl animal (Jax Stock#: 028866) to allow for the integration of a genetically encoded calcium indicator in all sensory neurons for calcium imaging. All procedures were approved by the CCHMC Institutional Animal Care and Use Committee in compliance with AALAC approved practices.

### Intranasal administration of *Alternaria Alternata*

Allergic esophageal inflammation was induced by intranasal administration of *Alternaria alternata* extract (Greer Laboratories, Inc.; 2 mg/dose in 30 ul of sterile phosphate-buffered saline [PBS]) three times a week for a total of 11-12 injections. Control mice were given 30 ul of PBS alone. Mice were sacrificed two days after their final allergen treatment for all analyses. Following saline perfusion, esophagi were fixed in 4% paraformaldehyde for histological or immunofluorescence analysis unless otherwise specified. All tissue processing and histological staining (H&E and MBP) slides were generated with the help of the CCHMC Pathology Core.

### Behavioral assays

Mice were tested for baseline behavior prior to any allergen administration. Mice were then analyzed again at day 7, 14, 21, and 26 prior to the intranasal allergen/saline administration for that day. On days 0, 7, 14, 21, and 26, two behavioral tasks were performed: assessment of mouse grimace scale (nonevoked) and analysis of mechanical withdrawal scoring to epigastric von Frey probes. Non-evoked pain-like behavior was measured by qualitatively assessing five mouse facial characteristics on a scale of zero to two, where zero is no change from normal in facial feature, 1 indicates moderate presence of changes, and 2 indicates obvious changes. Reference features were acquired from Rangford et al. (17). Assessments were made every 5 min for 30 mins total. Epigastric-directed withdrawal scores were measured following application of von Frey probes at two different strengths (1g and 4g). In this assay, the animal was lightly poked in the epigastric region with a von Frey probe and withdrawal response was scored on a scale of zero to three, where zero is no response, 1 indicates sudden retraction of their body, 2 indicates retraction and licking to the area, and 3 indicates jumping. This was repeated for a total of three trials for each von Frey probe for each animal, separated by at least 5 min between trials for recovery.

### Dissections

Prior to dissection of any tissue, mice were first anesthetized through an intramuscular injection of ketamine/xylazine solution (9-mg/mL ket-amine 1 0.9-mg/mL xylazine) and then cardiac perfused with ice-cold saline unless otherwise specified.

### Isolation, culture, and calcium imaging of DRG and VG cells

Mouse DRG and VG neurons from both sexes were isolated and cultured using previously published protocols with slight modification (62–64). Bilateral T1-T5 DRGs and bilateral VG were removed from Pirt-GCaMP6 mice unless otherwise specified. After removal, ganglia were transferred to 1 mL Ca2+/Mg2+-free Hank’s Balanced Salt Solution (HBSS). Ganglia were first incubated in 1.5 mL of Ca2+/Mg2+-free HBSS containing 1 mg L-cysteine (Sigma-Aldrich), and 60 U papain (Worthington) and incubated at 37C for 10 min. The suspension was then centrifuged, and the supernatant was removed and replaced with 1.5 mL Ca2+/Mg2+-free HBSS containing 12 collagenase type II (Worthington) and incubated at 37C for 10 min. After digestion, neurons were gently triturated, pelleted, and then resuspended in Ham’s F-12 Nutrient Mixture (Thermo Fischer), 100 U/mL penicillin plus 100 mg/mL streptomycin, and 10% heat-inactivated FBS (Sigma-Aldrich). Neurons were then plated on glass coverslips (Marienfeld) in 12-well plate, pre-coated with poly-L-lysine (Sigma-Aldrich) and laminin (Sigma-Aldrich) and cultured under a humidified atmosphere of 5% CO2 at 37C for 1 hr. Wells were flooded with 1mL of F-12 mixture and then incubated overnight for 18–24 hr before use. Prior to imaging, cultured mouse DRG and VG neurons were incubated in Kreb’s Henseleit Buffer (132 mM NaCl, 4.7 mM KCl, 2.0 mM CaCl2, 2.0 mM MgSO4, 21.8 mM NaHCO3, pH 7.45) at room temperature for 30 min before use.

Murine recombinant IL-4,(Peprotech and R&D Systems), capsaicin, adenosine triphosphate (ATP), and mustard oil *were used at* 300 nM, 1 uM, 150 nM, and 1 uM respectively, unless otherwise indicated. Where indicated, sensory neurons were also challenged with corresponding dilutions of control vehicle (0.1% BSA (Sigma-Aldrich) in PBS). Only sensory neurons that responded to a final challenge of 50 mM KCl were used in analyses. Fluorescence was recorded at 340 nm and 380 nm excitation wavelengths (F340, F380) using an inverted Nikon A1 microscope with NIS-Elements imaging software (Nikon Instruments). Calcium transients were measured for after stimulation with specified stimulus. dF/F0 were calculated where dF/F0 = (Fmax – F0)/F0 where Fmax: maximum intensity during stimulation and F0: the average intensity of immediately prior to stimulation. Data were then analyzed in Nikon Elements where cells were considered responsive if they demonstrated a change in fluorescence ratio (dF/F0) >1.

### RNA isolation and Real-Time PCR

#### Dorsal root and nodose ganglia

Using previously published procedures (34,65,66) and primers (34,67), all ganglia were dissected as described. Following dissection, ganglia were immediately frozen on dry ice until all dissections were completed (maximum of 4 hours). Samples were then used for RNA isolation using Qiagen RNeasy Mini kits according to the manufactures’ directions.

#### Esophagus

Following dissection, esophagi were placed in RNAlater solution (#AM7020) and kept at −80C until use. Samples were then used for the RNA isolation ZYMO Extraction Kit according to the manufactures’ directions. An Epoch microplate reader was used with the program Gen5 to assess quantification and purity.

#### Real-time PCR

cDNA was prepared from at least five hundred nanograms of total RNA by ProtoScript® First Strand cDNA Synthesis Kit following manufacture instructions. Samples were stored at −80 C° until needed for real-time PCR. To perform real-time RT polymerase chain reaction 20 ng of cDNA per well was used with SYBR-Green real-time PCR reactions (Applied Biosystems, Waltham, MA) and necessary primers on a StepOne real-time PCR System (Applied Biosystems). Relative expression was calculated by normalization of the Ct values of the targets to the Ct value of GAPDH for each well. Fold changes between conditions were converted to a percent change where 2-fold =100% change. MIQE guidelines were followed to ensure integrity.

### Bulk RNA-sequencing

Following RNA isolation as described above, all samples were sent to Novogene Bioinformatics Technology Co. (Novogene.com), where all bulk sequencing steps of RNA-seq were performed. All samples from all experiments were processed using the same pipeline for compatibility. The GRCm38 genome assembly was used as a genome and transcriptome track, with Mus musculus mm10 was used as the reference sequence. CLC Genomics Workbench version 24.0.2 was used for all analyses. Individual statistical tests and cutoffs for differential expression analyses are described in the manuscript.

### Immunofluorescence (IF)

#### Human esophagus

Formalin-fixed paraffin-embedded (FFPE) human esophageal biopsies were cut into 5 µM sections. Slides were deparaffinized with xylene, subjected to graded ethanol washes and subsequent antigen retrieval with antigen retrieval buffer under pressurized conditions, washed in PBS, and blocked in 10% donkey serum in PBS. Biopsies were incubated with primary antibodies (anti-BIII-tubulin (Biolegend 801201), anti-E Cadherin (R&D AF648), anti-EPX (MM25.82.2.1), and anti-tryptase (#369402, Biolegend))overnight at 4°C. The next day, the tissue was washed and stained with secondary antibodies and Hoechst 33342 for DNA staining. Images were taken on a Nikon A1R LUN-V inverted confocal microscope.

#### Murine esophagus

Prior to fixation, an open-book preparation was obtained by sectioning the esophagus longitudinally. Esophagi were fixed for 24 hours in 4% paraformaldehyde, then stored in 30% sucrose at 4°C. Samples were processed, cut into 15 µM frozen sections, embedded on microscope slides by CCHMC Pathology Core, and kept at −20C until use. Upon use, slides were thawed and rehydrated in PBS for 1 hour at room temperature. Slides were blocked in 10% donkey serum in PBS/0.2% Triton for 1 hour. They were then incubated with anti-BIII-tubulin (1:400) overnight at 4C. The next day, the samples were washed and stained with donkey-anti-rabbit secondary along with Hoechst 33342. After 2 hour incubation, washed samples with PBS then cover slipped until imaging with Nikon A1R LUN-V inverted confocal.

#### DRG/VG

For DRG extraction, after perfusion with ice-cold 0.9% saline solution, a laminectomy was performed to expose the spinal cord, and then, the T4 DRG and VG were isolated. After tissue dissection, ganglia were immersion fixed in 4% paraformaldehyde for 30 minutes before being embedded in 10% gelatin (in MQ water) and post-fixed overnight in 3% PFA. The gelatin blocks are then rinsed and placed in 20% sucrose (in 0.1M PB) overnight or About 45-um ganglia sections were cut using an HM 430 sliding microtome and placed in a 12-well plate where they were first rinsed in 0.1-M PBS. Slides/wells were then blocked in 0.01-M PBS with 4% donkey serum, 1% bovine serum albumin, and 0.01-M PBS with 0.1% Tween-20 for 1 hour. Then, the slides/wells were incubated in the above blocking buffer overnight at room temperature with gentle rocking with the primary antibodies as indicated by rabbit anti-TRPV1 (1:2000, Alomone), rabbit anti-TRPA1 (1:1000) or guinea pig anti-P2X3 (1:2000, Thermo Scientific). Next, slides/wells were washed in 0.01-M PBS and incubated in the same blocking buffer but with appropriate secondary antibodies (647 donkey anti-guinea pig, and 594 donkey anti-rabbit, Jackson Immunoresearch, West Grove, PA). The gelatin-embedded DRG sections in the wells were then placed carefully on gelatin-coated slides. The slides were then rinsed in 0.01-M PBS and coverslipped using Fluro media with DAPI to stain nuclei. Labeling of cells was identified and characterized on a Nikon A1 inverted confocal microscope with sequential scanning. Images were then compiled and prepared for publishing through NIS-elements imaging software.

For quantification, nonsequential sections of samples were counted to prevent any overlapping cells from being quantified more than once. Cells that were positive for one or more antibodies were counted separately, and each marker was quantified for each section. Counts were performed on 3 sections per single animal, and the average counts per condition were used for analysis. Positive cells were counted through ImageJ using each individual channel. Negative controls were performed by staining the slides with only secondary antibodies (ie, no primary) to confirm specificity.

### Quantification of nerve density

Tissue area measurements were first calculated using low-level whole tissue autofluorescence at 570 nm wavelength. Images of mouse and human esophageal biopsies were processed with Nikon Elements software. Once the intensity threshold was set, regions of interest (ROI) were automatically detected in Nikon Elements and the total area of each signal was quantified. BIII-tubulin signal was used to quantify myelinated nerve area in humans and mice; expression of the endogenous NaV1.8-Tdtomato reporter signal was used to quantify nociceptor nerve area in mice. Nerve area (BIII-tubulin or Tdtomato signal) was then divided by the size of the surveyed area and multiplied by 100 to calculate the innervation density. All investigators were blinded during quantifications. For human samples, at least 2 view fields per biopsy were imaged at 10x magnification and analyzed. Next, average densities were calculated using all images per patient. Every data point on a given graph corresponds to a single patient. For mouse samples, at least 2 images along the length of the esophagus and 2 consecutive cuts per mouse esophagus were imaged at 10x magnification and analyzed. Every data point on a given graph corresponds to a single image.

### Statistical analyses

Data was analyzed and graphed using Prism 10 software (GraphPad Software Inc.). All data were first checked for normality by Shapiro-Wilk to determine parametric or nonparametric tests. Individual tests used are described in the figure legends. Behavioral data of the same animal over time and between groups were analyzed using a two-way repeated measures (RM) ANOVA. In situations in which different animals were compared across multiple time points and groups, such as PCR data, a two-way ANOVA was used. In all experiments in which there is a potential for bias including behavior, innervation quantification, and IHC the investigator was blinded to the conditions. The data are shown as the mean ± standard error of the mean (SEM), unless otherwise indicated. Data from independent experiments are representative of at least two independent replicates or as pooled data. None of the data were excluded from statistical analyses, unless due to technical errors. Significance is regarded as: ****p < 0.0001, ***p < 0.001, **p < 0.01, *p < 0.05, and n.s. as not significant.

## References

1. Dellon ES. Eosinophilic esophagitis: Diagnostic tests and criteria. 2015

2. Bredenoord AJ, Patel K, Schoepfer AM, Dellon ES, Chehade M, Aceves SS, et al. Disease Burden and Unmet Need in Eosinophilic Esophagitis. 2022;117.

3. Safroneeva E, Straumann A, Schoepfer A. Latest Insights on the Relationship between Symptoms and Biologic Findings in Adults with Eosinophilic Esophagitis. 2021

4. Chehade M, Dellon ES, Spergel JM, Collins MH, Rothenberg ME, Pesek RD, et al. Dupilumab for Eosinophilic Esophagitis in Patients 1 to 11 Years of Age. N Engl J Med. 2024 Jun 27;390(24):2239–51.

5. Kliewer KL. Benralizumab for eosinophilic gastritis: a single-site, randomised, double-blind, placebo-controlled, phase 2 trial. 2023;8.

6. Oetjen LK. Sensory Neurons Co-opt Classical Immune Signaling Pathways to Mediate Chronic Itch.

7. Tamari M. Sensory neurons promote immune homeostasis in the lung. OPEN ACCESS.

8. Talbot S. Silencing Nociceptor Neurons Reduces Allergic Airway Inflammation.

9. Kim B. Neuroimmune interplay during type 2 inflammation: Symptoms, mechanisms, and therapeutic targets in atopic diseases. J ALLERGY CLIN IMMUNOL. 2024;153(4).

10. Yu M, Chang C, Undem BJ, Yu S. Capsaicin-Sensitive Vagal Afferent Nerve-Mediated Interoceptive Signals in the Esophagus. 2021;

11. Sengupta JN. An Overview of Esophageal Sensory Receptors. Am J Med. 2000;108(4A):87– 9.

12. Fass R. Sensory Testing of the Esophagus. J Clin Gastroenterol. 2004;38(8).

13. Mayo Clinic Staff. Mayo Clinic. 2019. Eosinophilic Esophagitis. Available from: https://www.mayoclinic.org/diseases-conditions/eosinophilic-esophagitis/symptoms-causes/syc-20372197

14. Lowenstein ED. Prox2 and Runx3 vagal sensory neurons regulate esophageal motility. OPEN ACCESS.

15. Mishra A. Significance of Mouse Models in Dissecting the Mechanism of Human Eosinophilic Gastrointestinal Diseases (EGID). 2015;

16. Mishra A, Hogan SP, Brandt EB, Rothenberg ME. An etiological role for aeroallergens and eosinophils in experimental esophagitis. J Clin Invest. 2001;107(1).

17. Langford D, Bailey A, Chanda M. Coding of facial expressions of pain in the laboratory mouse. Nat Methods. 2010;(7):447–9.

18. Dourson AJ. Macrophage memories of early-life injury drive neonatal nociceptive priming. OPEN ACCESS.

19. Gunasekaran T, Prabhakar G, Schwartz A, Gorla K, Gupta S, Berman J. Eosinophilic Esophagitis in Children and Adolescents with Abdominal Pain: Comparison with EoE-Dysphagia and Functional Abdominal Pain. Can J Gastroenterol Hepatol.

20. Deng L. Sensory neurons: An integrated component of innate immunity.

21. Kupari J. An Atlas of Vagal Sensory Neurons and Their Molecular Specialization.

22. Jung M. Cross-species transcriptomic atlas of dorsal root ganglia reveals species-specific programs for sensory function. Nat Commun. 2023;

23. Fountain SJ. Primitive ATP-activated P2X receptors: discovery, function and pharmacology. Front Cell Neurosci.

24. Yang F. Understand spiciness: mechanism of TRPV1 channel activation by capsaicin.

25. Bandwell M, Story G, Hwang SW, Viswanath V, Eid S, Parapoutian A. Noxious Cold Ion Channel TRPA1 Is Activated by Pungent Compounds and Bradykinin. Neuron. 2004 Mar 25;41:849–57.

26. Iadarola MJ. Comparative Analysis of Dorsal Root, Nodose and Sympathetic Ganglia for the Development of New Analgesics. Front Neurosci. 2020;14.

27. Jassal B, Matthews L, Viteri G, Gong C, Lorente P, Fabregat A, et al. The reactome pathway knowledgebase. Nucleic Acids Res. 2020 Jan 8;48(D1):D498–503.

28. Gautron L, Sakata I, Udit S, Zigman JM, Wood JN, Elmquist JK. Genetic Tracing of Nav1.8-Expressing Vagal Afferents in the Mouse. 2012;

29. Jiang Y, Dong H, Eckmann L, Hanson EM, Ihn KC, Mittal RK. Visualizing the enteric nervous system using genetically engineered double reporter mice: Comparison with immunofluorescence. PLOS ONE. 2017;

30. Foster EL, Simpson EL, Fredrikson LJ, Lee JJ, Lee NA, Fryer AD, et al. Eosinophils Increase Neuron Branching in Human and Murine Skin and In Vitro. Tsokos GC, editor. PLoS ONE. 2011 Jul 21;6(7):e22029.

31. Bao C, Abraham SN. Mast cell–sensory neuron crosstalk in allergic diseases. J Allergy Clin Immunol. 2024 Apr;153(4):939–53.

32. Zhang S, Shoda T, Aceves SS, Arva NC, Chehade M, Collins MH, et al. Mast cell-pain connection in eosinophilic esophagitis. Allergy. 2022 Jun;77(6):1895–9.

33. Ren K. Role of interleukin-1β during pain and inflammation. 2011;

34. Ross XJL, Queme XLF, Cohen XER, Green XKJ, Lu P, Shank AT, et al. Muscle IL1␤ Drives Ischemic Myalgia via ASIC3-Mediated Sensory Neuron Sensitization.

35. Lebold KM, Drake MG, Hales-Beck LB, Fryer AD, Jacoby DB. IL-5 Exposure In Utero Increases Lung Nerve Density and Airway Reactivity in Adult Offspring. 2020;62(4).

36. Blanchard C, Mingler MK, Vicario M, Pablo J, Wu YY, Lu TX, et al. IL-13 involvement in eosinophilic esophagitis: Transcriptome analysis and reversibility with glucocorticoids. J ALLERGY CLIN IMMUNOL. 2007;120(6).

37. Mishra A, Rothenberg ME. Intratracheal IL-13 Induces Eosinophilic Esophagitis by an IL-5, Eotaxin-1, and STAT6-Dependent Mechanism. 2003;125(5).

38. Oetjen LK, Mack MR, Feng J, Whelan TM, Niu H, Guo CJ, et al. Sensory Neurons Co-opt Classical Immune Signaling Pathways to Mediate Chronic Itch. Cell. 2017 Sep;171(1):217–228.e13.

39. Daines JM, Schellhardt L, Wood MD. The Role of the IL-4 Signaling Pathway in Traumatic Nerve Injuries. Neurorehabil Neural Repair. 2021 May;35(5):431–43.

40. Pan D, Schellhardt L, Acevedo-Cintron JA, Hunter D, Snyder-Warwick AK, Mackinnon SE, et al. IL-4 expressing cells are recruited to nerve after injury and promote regeneration. Exp Neurol. 2022 Jan;347:113909.

41. Safroneeva E, Straumann A, Schoepfer AM. Latest Insights on the Relationship Between Symptoms and Biologic Findings in Adults with Eosinophilic Esophagitis. Gastrointest Endosc Clin N Am. 2018 Jan;28(1):35–45.

42. Dourson AJ, Ford ZK, Green KJ, McCrossan CE, Hofmann MC, Hudgins RC, et al. Early Life Nociception is Influenced by Peripheral Growth Hormone Signaling.

43. Sullivan K, Kim B, Feldman E. Insulin-Like Growth Factors in the Peripheral Nervous System. Endocrinology. 2008 Dec;149(12):5963–71.

44. Aghanoori MR, Agarwal P, Gauvin E, Nagalingam RS, Bonomo R, Yathindranath V, et al. CEBPβ regulation of endogenous IGF-1 in adult sensory neurons can be mobilized to overcome diabetes-induced deficits in bioenergetics and axonal outgrowth. Cell Mol Life Sci. 2022 Apr;79(4):193.

45. Lu TX, Lim EJ, Besse JA, Itskovich S, Plassard AJ, Fulkerson PC, et al. miR-223 Deficiency Increases Eosinophil Progenitor Proliferation. J Immunol. 2013 Feb 15;190(4):1576–82.

46. Veraldi KL, Gibson BT, Yasuoka H, Myerburg MM, Kelly EA, Balzar S, et al. Role of Insulin-like Growth Factor Binding Protein-3 in Allergic Airway Remodeling. Am J Respir Crit Care Med. 2009 Oct 1;180(7):611–7.

47. Rochman M, Wen T, Kotliar M, Dexheimer PJ, Ben-Baruch Morgenstern N, Caldwell JM, et al. Single-cell RNA-Seq of human esophageal epithelium in homeostasis and allergic inflammation. JCI Insight. 2022 Jun 8;7(11):e159093.

48. Qi L, Huang C, Wu X, Tao Y, Yan J, Shi T, et al. Hierarchical Specification of Pruriceptors by Runt-Domain Transcription Factor Runx1. J Neurosci. 2017 May 31;37(22):5549–61.

49. Elias LJ, Succi IK, Schaffler MD, Foster W, Gradwell MA, Bohic M, et al. Touch neurons underlying dopaminergic pleasurable touch and sexual receptivity. Cell. 2023 Feb;186(3):577–590.e16.

50. Rochman M. Epithelial origin of eosinophilic esophagitis. J ALLERGY CLIN IMMUNOL. 2018;142(1).

51. O’Shea KM, Aceves SS, Dellon ES, Gupta SK, Spergel JM, Furuta GT, et al. Pathophysiology of Eosinophilic Esophagitis. 2019;

52. Underwood B. Breaking down the complex pathophysiology of eosinophilic esophagitis. Ann Allergy Asthma Immunol. 2023;

53. Drake MG, Scott GD, Blum ED, Lebold KM, Nie Z, Lee JJ, et al. Eosinophils increase airway sensory nerve density in mice and in human asthma. 2019;

54. Collison AM, Sokulsky LA, Nightingale S, Percival E, LeFevre A, Meredith J, et al. *In vivo* targeting of miR-223 in experimental eosinophilic oesophagitis. Clin Transl Immunol. 2020 Jan;9(11):e1210.

55. Dellon ES, Rothenberg ME, Collins MH, Hirano I, Chehade M, Bredenoord AJ, et al. Dupilumab in Adults and Adolescents with Eosinophilic Esophagitis. N Engl J Med. 2022;

56. Lessmann E, Grochowy G, Weingarten L, Giesemann T, Aktories K, Huber M. Insulin and insulin-like growth factor-1 promote mast cell survival via activation of the phosphatidylinositol-3-kinase pathways. Exp Hematol. 2006;34:1532–41.

57. Alfaro-Arnedo E, López IP, Piñeiro-Hermida S, Ucero ÁC, González-Barcala FJ, Salgado FJ, et al. IGF1R as a Potential Pharmacological Target in Allergic Asthma. 2021;

58. Spadaro O, Camell CD, Bosurgi L, Nguyen KY, Youm YH, Rothlin CV, et al. IGF1 Shapes Macrophage Activation in Response to Immunometabolic Challenge. Cell Rep. 2017 Apr;19(2):225–34.

59. Granja MG, Gomes Braga LE, Monteiro De Oliveira R, De Mello Silva E, Gonçalves-de-Albuquerque CF, Silva AR, et al. IGF-1 and IGF-1R modulate the effects of IL-4 on retinal ganglion cells survival: The involvement of M1 muscarinic receptor. Biochem Biophys Res Commun. 2019 Oct;519(1):53–60.

60. Chu C. Neuro-immune Interactions in the Tissues.

61. Nagashima H. Neuropeptide CGRP Limits Group 2 Innate Lymphoid Cell Responses and Constrains Type 2 Inflammation.

62. Molliver DC, Rau KK, McIlwrath SL, Jankowski MP, Koerber HR. The ADP receptor P2Y1 is necessary for normal thermal sensitivity in cutaneous polymodal nociceptors. Mol Pain. 2011 Feb 10;7:13.

63. Malin SA, Davis BM, Molliver DC. Production of dissociated sensory neuron cultures and considerations for their use in studying neuronal function and plasticity. Nat Protoc. 2007;2(1):152–60.

64. Oetjen LK. Sensory Neurons Co-opt Classical Immune Signaling Pathways to Mediate Chronic Itch.

65. Dourson AJ, Fadaka AO, Warshak AM, Paranjpe A, Weinhaus B, Queme LF, et al. Macrophage memories of early-life injury drive neonatal nociceptive priming. Cell Rep. 2024 May;43(5):114129.

66. Quijas MM, Queme LF, Woodke ST, Weyler AA, Buesing D, Butterfield A, et al. Sex-specific role of RNA-binding protein, pAUF1, on prolonged hypersensitivity after repetitive ischemia with reperfusion injury. Pain. 2024 Oct 8;

67. Talbot S. Silencing Nociceptor Neurons Reduces Allergic Airway Inflammation.

