## Supplemental Figure Legends for "Identification of a Neuroimmune Circuit that Regulates Allergic Inflammation in the Esophagus"

**Supplementary figures:**

**Figure S1. Repeated allergen challenge induces EoE-like changes in murine esophagus. Related to Figure 1**

(A) Representative esophagus histology images of saline-control mice and mice challenged with intranasal allergen, obtained by H&E stain depicting epithelial disruption (*left*) and basal cell hyperplasia (*middle*), and eosinophilic major basic protein (MBP) stain (*right*) to visualize esophageal eosinophil infiltrate.

(B-C) Epigastric withdrawal thresholds in the von Frey test with 1g (B) and 4g (C) filaments, in saline control (black) and allergen treated (red) mice prior to intranasal allergen administration . Data are mean ± SEM of n = 6 mice per group. Two-way ANOVA with Sidaks multiple comparisons.

**Figure S2. Calcium imaging reveals little changes in VG response properties after capsaicin, ATP, and mustard oil exposure in allergic vs saline control mice. Related to Figure 1.**

(A-B) Quantification of immunohistochemistry staining of TRPV1+, P2X3+, and TRPA1+ in saline control vs allergen treated DRGs (A) and VGs (B).

(C-D) ATP responsive (C) and capsaicin (D) responsive VG neurons as a percentage of total KCl-responsive neurons (*left*), fluorescence intensity changes (*middle*), and representative calcium transients from sensory neurons of PBS and ALT treated mice in response to capsaicin (1 um) or ATP (150 um) stimulation. The arrow indicates when the stimulus was added. Each dot represents one cell. (*right*)

(E-F) Mustard oil (1um) responsive VG neurons (E) and DRG neurons (F) as a percentage of total KCl-responsive neurons (*left*) and fluorescence intensity changes (*right*). VG n>150 cells and DRG n>25 neurons from at least 3 Pirt-GCAMP6 mice per stimuli.

Percent responders: chi squared test, fluorescent intensity changes: unpaired T-test; VG n>150 from ≥3 Pirt-GCAMP6 mice. *p<0.05, ****p<0.0001; Data shown as mean +/- SEM.

**Figure S3. Whole DRG transcriptomic analysis separates allergen treated vs saline controls into two distinct groups. Related to Figure 2.**

(A) Heat map of 108 overlapping genes that are differentially expressed between allergen treated (red) and saline control (black) mice. FC threshold of 1.5 and p-value <0.05 between two experiments was applied; 15 downregulated and 93 upregulated.

**Figure S4. Whole murine esophagus reveals detailed innervation patterns of nociceptive and myelinated nerves. Related to Figure 3 and 4.**

(A) Video of whole esophagus from NaV1.8-Cre R26CAG-floxStop-tdTomato mice. Shown are NaV1.8+ nerves (red), basal epithelium (blue), BIII-tubulin (white) and muscle autofluorescence (green)

(B) Section of whole mouse esophagus from NaV1.8-Cre R26CAG-floxStop-tdTomato mice, showing intricate myenteric plexus formed from BIII-tubulin nerves (green)

(C) Section of whole mouse esophagus from NaV1.8-Cre R26CAG-floxStop-tdTomato mice, showing cluster of neurons forming the myenteric plexus with overlapping BIII-tubulin nerves (green) and NaV1.8 (red) nerves (purple).

(D) Schematic delineating the separation of esophageal layers for nerve density quantification. Related to Figure 4

**Figure S5. Human esophageal biopsies reveal distinct esophageal innervation during active EoE. Related to Figure 5.**

(A-B) H&E staining of human 35-week-old embryo (A) and 34-year-old donor (B) biopsies. Lumen is represented by the red asterisk.

(C) Representative immunofluorescence images of E-cadherin (green), DAPI (blue) and BIII-tubulin (purple) in esophageal biopsies from one healthy control and one active EoE patient. Positive BIII-tubulin nerve signal is shown in green on right images.

(D) Eosinophil levels in healthy control and remission patients (n=20) compared with Active EoE patients (n=38). Mann-Whittney test, ****p<0.0001; Each dot represents one patient.

(E) Correlation between BIII-tubulin innervation density and eosinophil levels per patient. Simple linear regression analysis on all patients. p=0.1627, R squared=03513

**Figure S6. Overnight IL-4 incubation does not alter DRG responsiveness to capsaicin and ATP. Related to Figure 6.**

(A-B) Capsaicin responsive (A) and ATP responsive (B) DRG neurons shown in color as a percentage of total KCl-responsive neurons shown in black (*top*) and fluorescence intensity changes (*bottom*). Cells are from DRG sensory neurons incubated overnight in regular media (gray) or IL-4 media (orange) in response to capsaicin (1 um) or ATP (150 um) stimulation. Percent responders: chi squared test, fluorescent intensity changes: unpaired T-test; DRG n>90 cells from ≥3 Pirt-GCAMP6 mice. Data shown as mean +/- SEM.

**Figure S7. Repeated allergen challenge alters esophageal inflammatory environment differently in WT compared to IL-4R-/-$^{NaV1.8}$ mice. Related to Figure 7.**

(A) The number of genes that exhibit the indicated minimum fold difference in mice given Alternaria (Allergic) versus saline controls is graphed (*left).* Genes with the highest fold change are listed (*right*).

(B) Venn-diagram demonstrating the number of differentially expressed genes in allergen-treated versus saline control mice (*right*) and allergen-treated WT versus allergen-treated IL-4R$^{NaV1.8}$ mice (*left*). DEGs that fit threshold criteria in both groups are shown in the center; 6 downregulated and 25 upregulated.

(C) Comparison of Alox15, Ccr3, and Rnase2 expression by qRT-PCR in whole mouse esophagus compared between saline control WT (dark gray), saline control IL-4R$^{NaV1.8}$ (light gray), allergen-treated WT (red) and allergen-treated IL-4R-/-$^{NaV1.8}$ (green). n > 3 mice per group. Kruskal-Wallis ANOVA test; statistics between red and green are the only ones shown *p<0.05, **p<0.005. Data shown as mean +/- range.

(D) Representative H&E staining of murine esophagus from IL-4R-/-$^{NaV1.8}$ (D) and WT (E) mice. Black arrow indicates epithelial cell disruption, and the yellow bracket showcases basal cell hyperplasia in the WT allergen-treated mouse.

**Figure S8. IL-4R-/-$^{NaV1.8}$ reduces DRG gene expression changes seen in following repeated allergen challenges. Related to Figure 8.**

(A) Venn-diagram demonstrating the number of differentially expressed genes in allergen-treated IL-4R-/-$^{NaV1}$ versus allergen-treated WT mice (*right*) and allergen-treated WT versus saline control WT mice (*left*). DEGs that fit threshold criteria in both groups are shown in the center.

(B) Pairwise comparison between the Log2 fold change of the 56 overlapping differentially expressed genes in each comparison (A). Green dots represent allergen-treated IL-4R-/-$^{NaV1}$ versus allergen-treated WT comparison and red dots represent allergen-treated WT versus saline control WT mice.

(C) List of the 56 overlapping differentially expressed genes and the corresponding numerical log2 fold change value between each group.
