## Supplemental Figures for "Identification of a Neuroimmune Circuit that Regulates Allergic Inflammation in the Esophagus"

### Slide 1
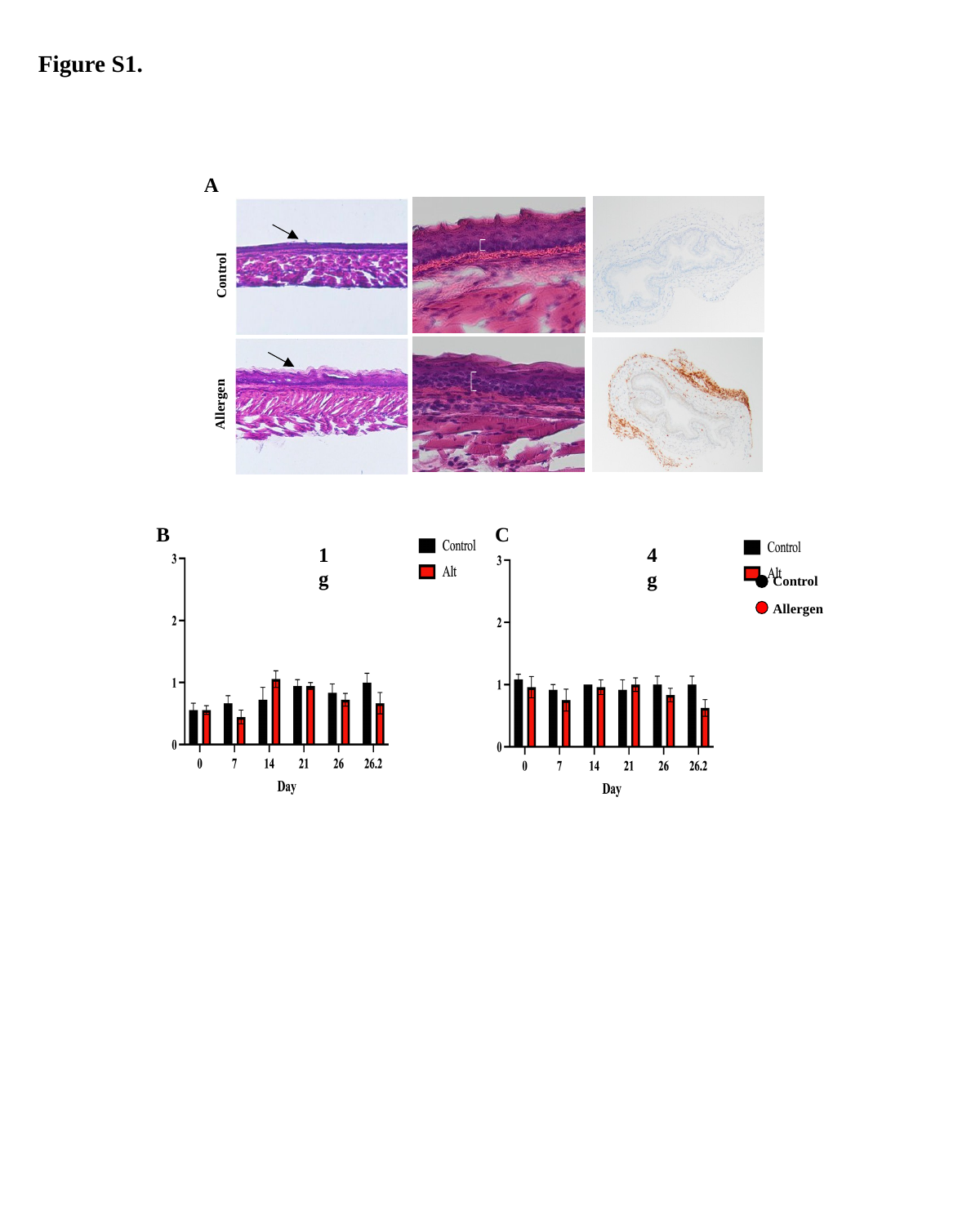

Figure S1.
A
Control
Allergen
B
C
1g
4g
Control
Allergen

### Slide 2
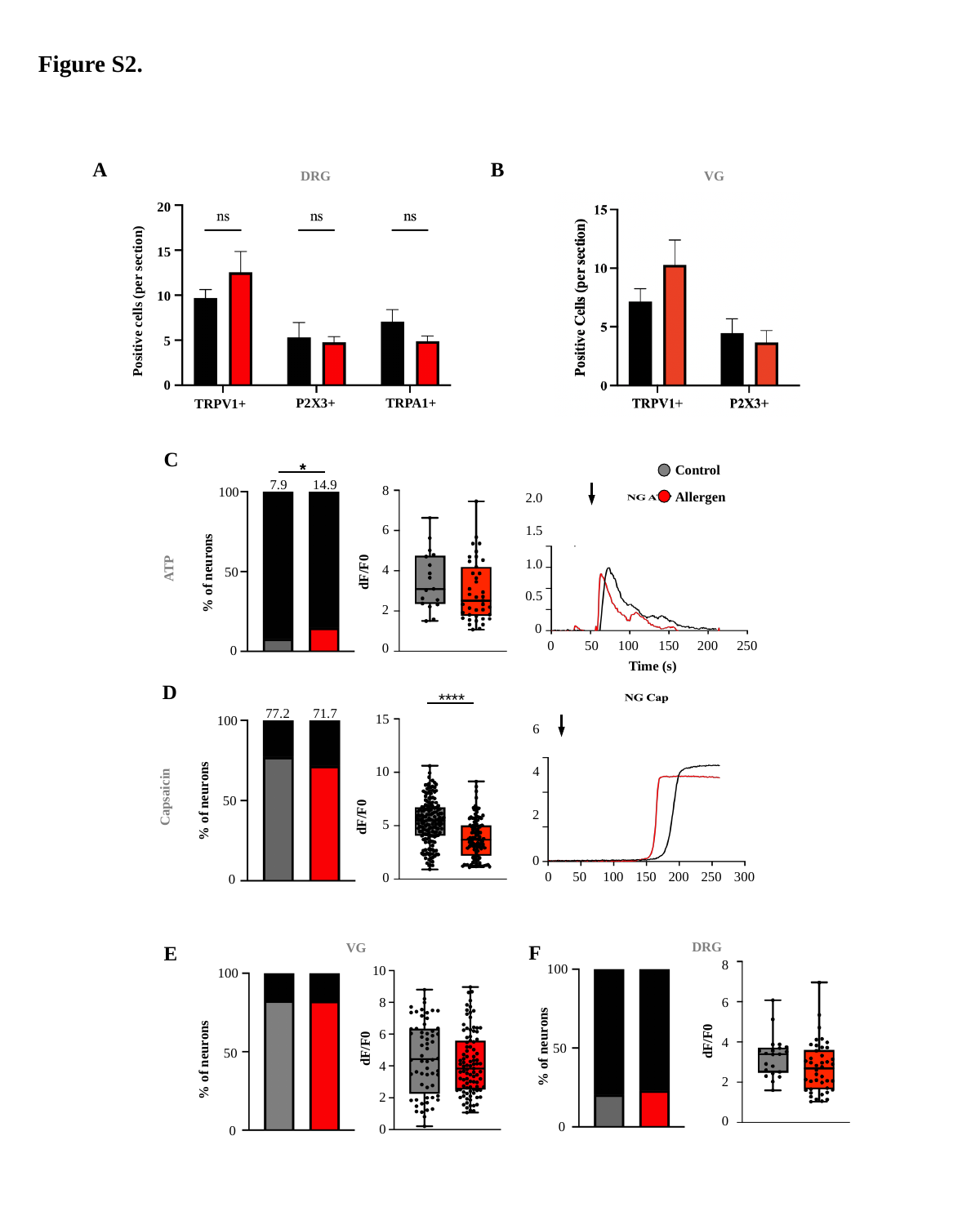

Figure S2.
A
B
DRG
VG
20
15
10
Positive cells (per section)
5
0
P2X3+
TRPA1+
TRPV1+
C
*
Control
7.9
14.9
8
6
4
2
0
100
2.0
1.5
1.0
0.5
0
0
50
100
150
200
250
Time (s)
Allergen
ATP
50
dF/F0
% of neurons
0
D
****
15
10
5
0
6
4
2
0
0
50
100
150
200
250
300
77.2
71.7
100
Capsaicin
50
% of neurons
dF/F0
0
DRG
VG
F
E
10
8
6
4
2
0
8
6
4
2
0
100
50
0
100
dF/F0
% of neurons
dF/F0
50
% of neurons
0

### Slide 3
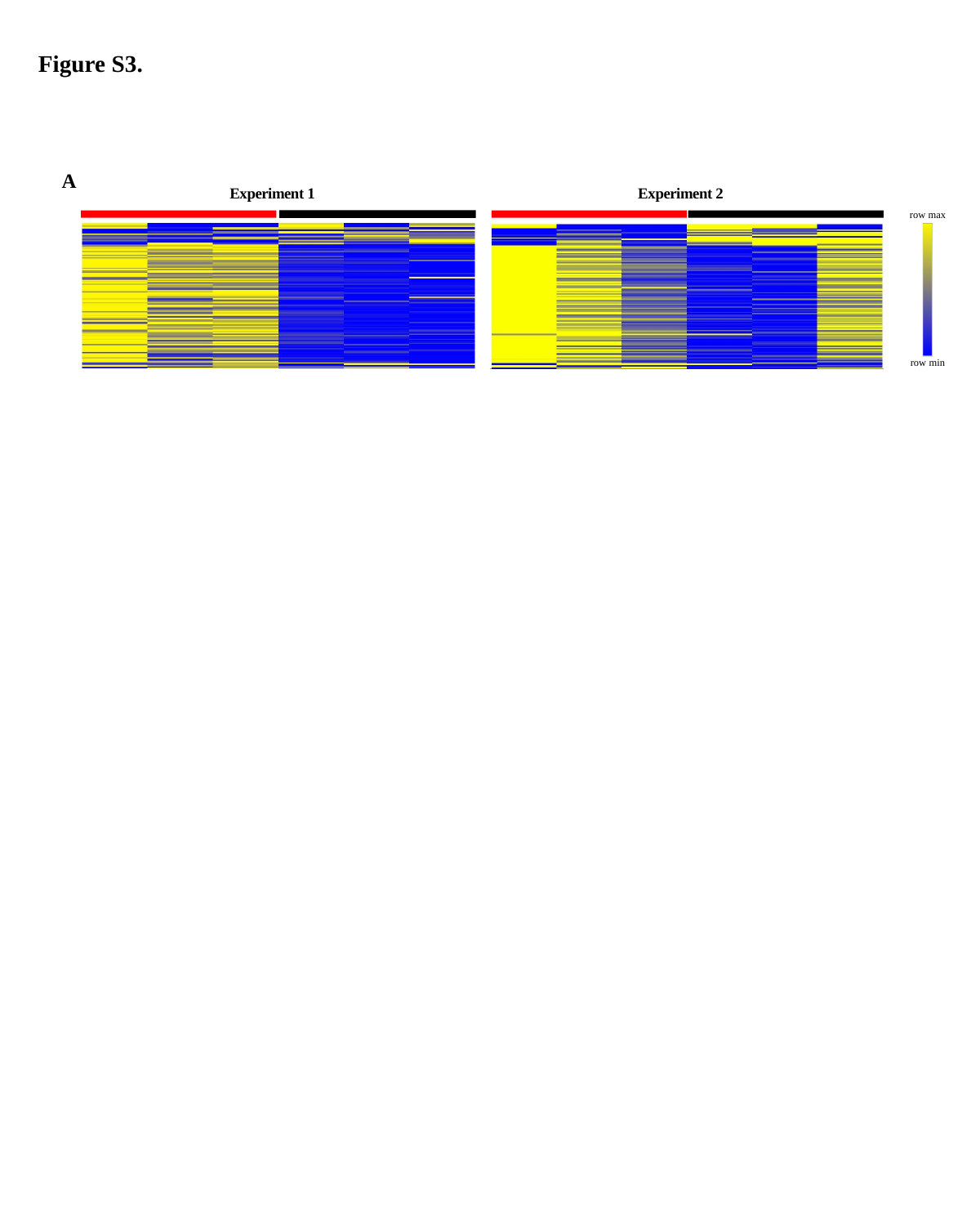

Figure S3.
A
Experiment 2
Experiment 1
row max
row min

### Slide 4
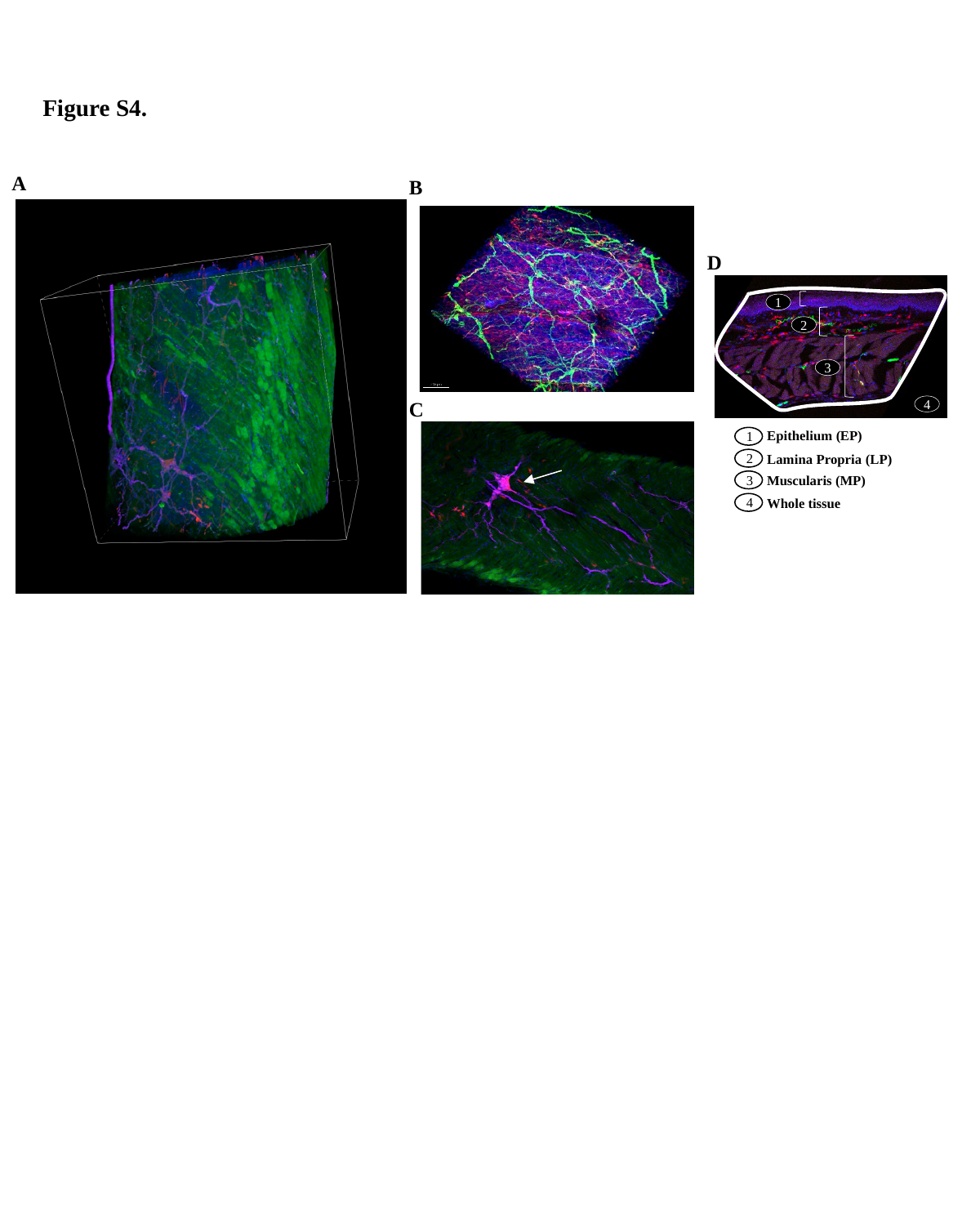

Figure S4.
A
B
D
1
2
3
C
4
Epithelium (EP)
1
2
Lamina Propria (LP)
3
Muscularis (MP)
4
Whole tissue

### Slide 5
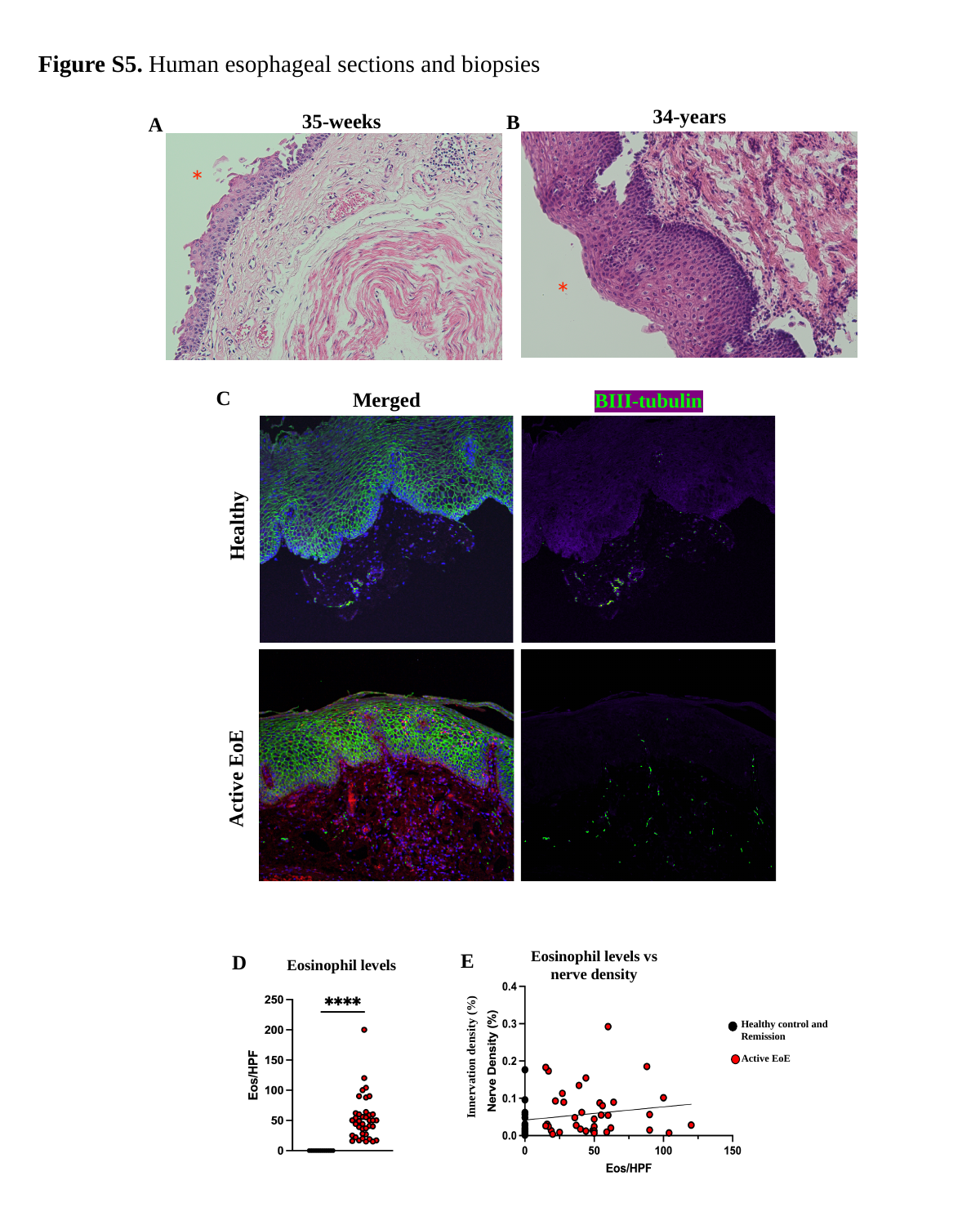

Figure S5. Human esophageal sections and biopsies
34-years
B
A
35-weeks
*
*
C
Merged
BIII-tubulin
Healthy
Active EoE
Eosinophil levels vs nerve density
E
D
Eosinophil levels
Innervation density (%)
Healthy control and Remission
Active EoE

### Slide 6
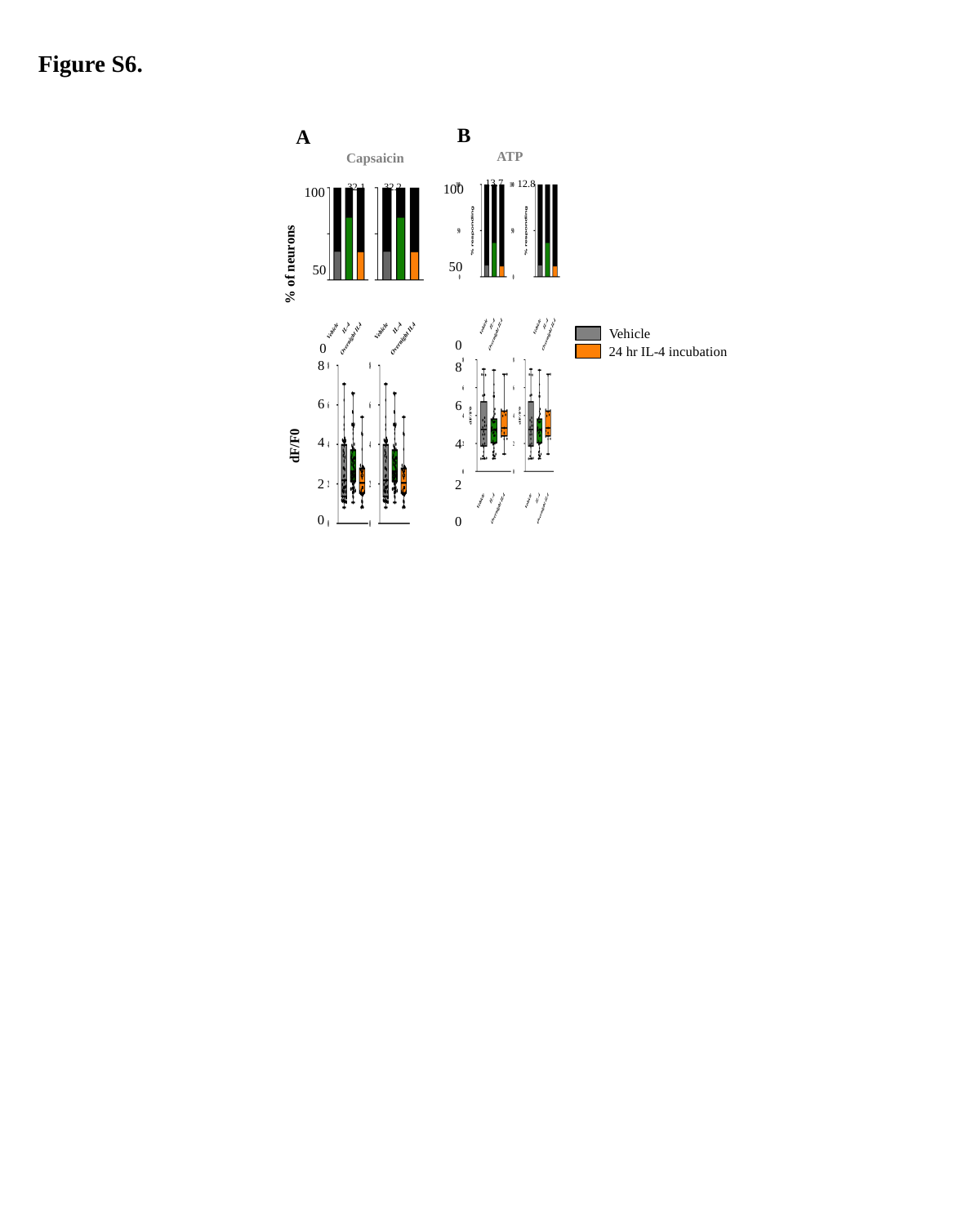

Figure S6.
B
A
ATP
Capsaicin
13.7
12.8
100
32.1
32.2
100
% of neurons
50
50
Vehicle
0
0
24 hr IL-4 incubation
8
6
4
2
0
8
6
4
2
0
dF/F0

### Slide 7
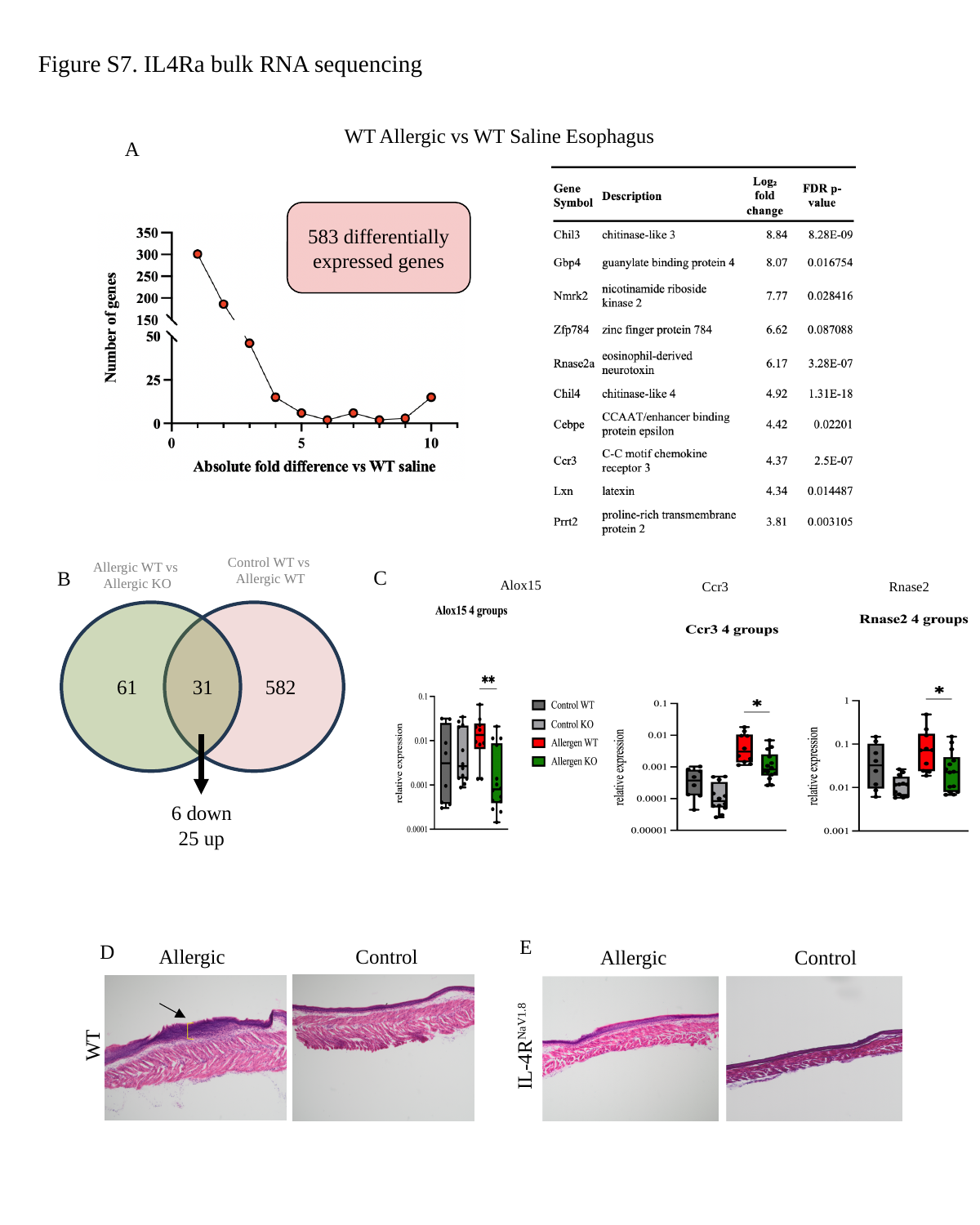

Figure S7. IL4Ra bulk RNA sequencing
WT Allergic vs WT Saline Esophagus
A
583 differentially expressed genes
582
Control WT vs
Allergic WT
6 down
25 up
31
Allergic WT vs
Allergic KO
61
C
B
Alox15
Ccr3
Rnase2
E
D
Allergic
Control
Allergic
Control
IL-4R^NaV1.8^
WT

### Slide 8
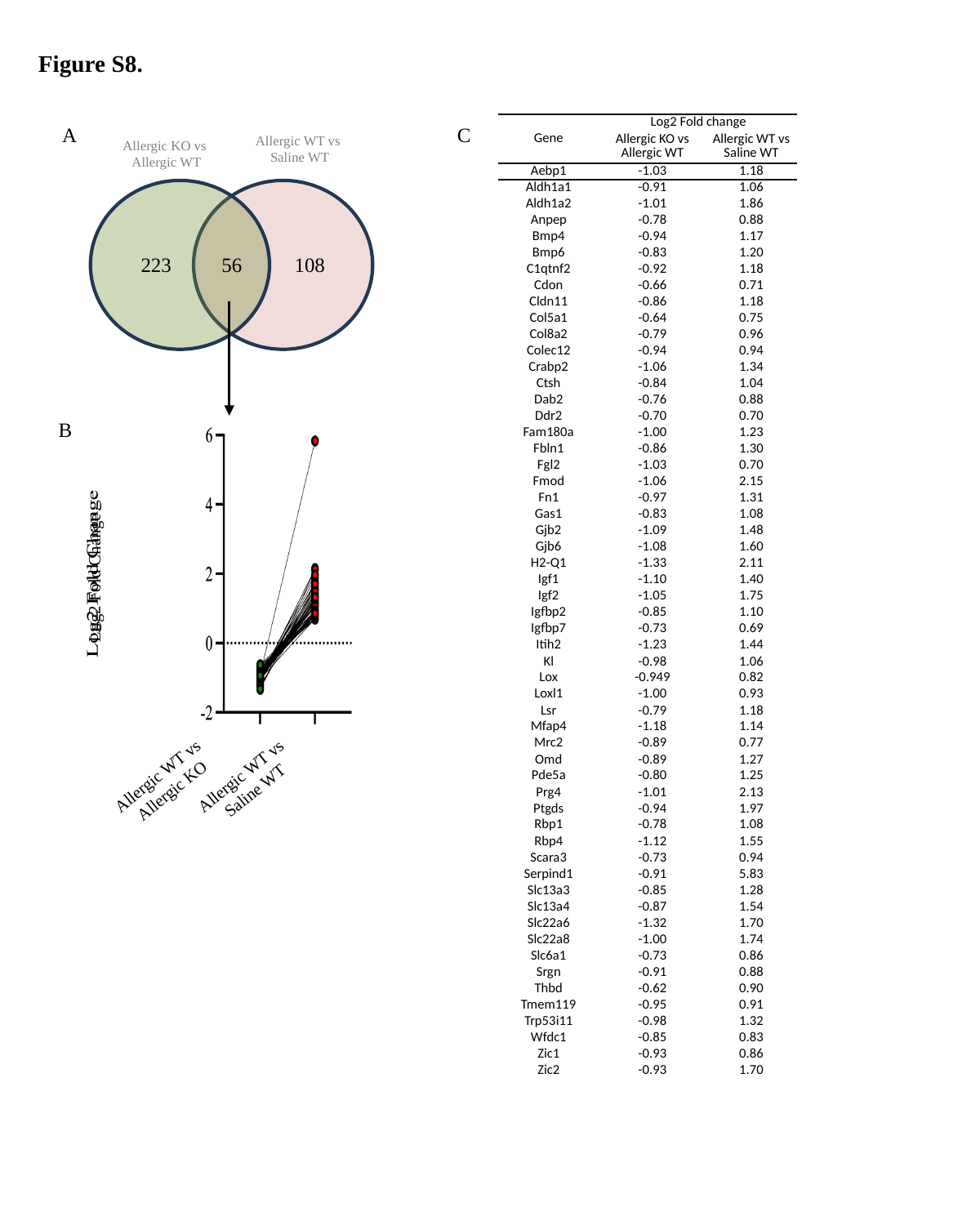

Figure S8.
108
Allergic WT vs
Saline WT
56
Allergic KO vs
Allergic WT
223
| Gene | Log2 Fold change | |
| --- | --- | --- |
| | Allergic KO vs Allergic WT | Allergic WT vs Saline WT |
| Aebp1 | -1.03 | 1.18 |
| Aldh1a1 | -0.91 | 1.06 |
| Aldh1a2 | -1.01 | 1.86 |
| Anpep | -0.78 | 0.88 |
| Bmp4 | -0.94 | 1.17 |
| Bmp6 | -0.83 | 1.20 |
| C1qtnf2 | -0.92 | 1.18 |
| Cdon | -0.66 | 0.71 |
| Cldn11 | -0.86 | 1.18 |
| Col5a1 | -0.64 | 0.75 |
| Col8a2 | -0.79 | 0.96 |
| Colec12 | -0.94 | 0.94 |
| Crabp2 | -1.06 | 1.34 |
| Ctsh | -0.84 | 1.04 |
| Dab2 | -0.76 | 0.88 |
| Ddr2 | -0.70 | 0.70 |
| Fam180a | -1.00 | 1.23 |
| Fbln1 | -0.86 | 1.30 |
| Fgl2 | -1.03 | 0.70 |
| Fmod | -1.06 | 2.15 |
| Fn1 | -0.97 | 1.31 |
| Gas1 | -0.83 | 1.08 |
| Gjb2 | -1.09 | 1.48 |
| Gjb6 | -1.08 | 1.60 |
| H2-Q1 | -1.33 | 2.11 |
| Igf1 | -1.10 | 1.40 |
| Igf2 | -1.05 | 1.75 |
| Igfbp2 | -0.85 | 1.10 |
| Igfbp7 | -0.73 | 0.69 |
| Itih2 | -1.23 | 1.44 |
| Kl | -0.98 | 1.06 |
| Lox | -0.949 | 0.82 |
| Loxl1 | -1.00 | 0.93 |
| Lsr | -0.79 | 1.18 |
| Mfap4 | -1.18 | 1.14 |
| Mrc2 | -0.89 | 0.77 |
| Omd | -0.89 | 1.27 |
| Pde5a | -0.80 | 1.25 |
| Prg4 | -1.01 | 2.13 |
| Ptgds | -0.94 | 1.97 |
| Rbp1 | -0.78 | 1.08 |
| Rbp4 | -1.12 | 1.55 |
| Scara3 | -0.73 | 0.94 |
| Serpind1 | -0.91 | 5.83 |
| Slc13a3 | -0.85 | 1.28 |
| Slc13a4 | -0.87 | 1.54 |
| Slc22a6 | -1.32 | 1.70 |
| Slc22a8 | -1.00 | 1.74 |
| Slc6a1 | -0.73 | 0.86 |
| Srgn | -0.91 | 0.88 |
| Thbd | -0.62 | 0.90 |
| Tmem119 | -0.95 | 0.91 |
| Trp53i11 | -0.98 | 1.32 |
| Wfdc1 | -0.85 | 0.83 |
| Zic1 | -0.93 | 0.86 |
| Zic2 | -0.93 | 1.70 |
A
C
B
Log2 Fold Change
Allergic WT vs
Allergic KO
Allergic WT vs
Saline WT
